# TOR kinase controls shoot development by translational repression of cytokinin catabolic enzymes

**DOI:** 10.1101/2021.07.29.454319

**Authors:** Denis Janocha, Anne Pfeiffer, Yihan Dong, Ondřej Novák, Miroslav Strnad, Isabel Bartrina, Lyuba A Ryabova, Tomas Werner, Jan U. Lohmann

## Abstract

Plants continuously adjust their developmental program including organ initiation and growth in accordance with endogenous and environmental signals. This plasticity requires that a diversity of signaling pathways acts in concert to modulate stem cell activity. We have shown previously that the TOR kinase network integrates metabolic- and light signals and controls expression of *WUSCHEL*, a transcriptional master regulator of stem cells in the shoot apical meristem. However, the mechanism linking TOR activity with the *WUSCHEL* promoter remained unresolved. Here we demonstrate that TOR regulates the accumulation of *trans*-zeatin, the cytokinin species mainly responsible for shoot development. Importantly, we identify translational repression of RNAs encoding cytokinin degrading CYTOKININ OXIDASES/DEHYDROGENASE enzymes by TOR as an underlying mechanism. Employing this system, plants can quickly adjust stem cell activity and developmental programs in response to changes in their environment.

## Introduction

Due to their sessile lifestyle, plants depend on their ability to dynamically adjust organ development and growth rates in response to variable environmental conditions. Therefore, they can integrate diverse local and systemic signals to adjust stem cell behavior accordingly. Stem cells of the shoot apical meristem (SAM) are controlled by the activity of the homeodomain transcription factor WUSCHEL (WUS) (Bäurle & Laux, 2005; Daum et al., 2014; Laux et al., 1996; Mayer et al., 1998; Schoof et al., 2000) and previous studies have shown that *WUS* expression is modulated in response to environmental signals such as nitrate availability or the current light- and energy regime (Janocha & Lohmann, 2018; Landrein et al., 2018; Li et al., 2017; Pfeiffer et al., 2016). In the absence of a neuronal system, plants make use of complex hormone signaling systems for long distance information relay. Among those, cytokinins (CKs) and particularly *trans*-zeatin (*tZ*), are involved in nitrate and light signaling from root and leaves to the SAM, respectively (Landrein et al., 2018; Yoshida et al., 2011). CKs in turn promote *WUS* expression, thus tuning stem cell number in response to the environment (Buechel et al., 2010; Gordon et al., 2009; Kiba et al., 2013; Landrein et al., 2018; Osugi et al., 2017).

CK homeostasis is controlled by tightly regulated biosynthesis and degradation systems (Heyl et al., 2018), with ISOPENTYLTRANSFERASES (IPTs) catalyzing the first step in the anabolic pathway. IPTs synthesize isopentyladenine-5’-mono-, di- and tri-phosphates (*i*PRnP) precursors, which can be hydroxylated by CYP735A1/2 to obtain *trans*-zeatin ribosides (*t*ZRs). Both precursors are then converted by LONELY GUY (LOGs) enzymes to obtain the active nucleobases isopentenyladenine (*i*P) and *tZ* (Kuroha et al., 2009). CK degradation in *Arabidopsis* is mediated by seven cytokinin oxidase/dehydrogenase (CKXs) enzymes, which are characterized by distinct tissue specific expression making them important players to locally and globally control CK signaling (Bartrina et al., 2011; Holst et al., 2011; Köllmer et al., 2014; X. Wang et al., 2020; Werner et al., 2003). Intracellular CK signal transduction is based on a two-component phospho-relay (Kieber & Schaller, 2014). The receptors are *ARABIDOPSIS HISTIDINE KINASES* (*AHK*s) and reside on the plasma membrane and the endoplasmic reticulum (Antoniadi et al., 2020). Upon CK binding, AHKs first auto-phosphorylate and then pass on the phosphoryl group to *ARABIDOPSIS HISTIDINE PHOSPHOTRANSFER PROTEINS* (*AHP*s) in an evolutionary conserved phospho-relay. AHPs continuously shuttle between cytoplasm and nucleus where they interact with and activate type-B *ARABIDOPSIS RESPONSE REGULATORS* (*ARR*s) transcription factors by phosphorylation (Suzuki et al., 1998, 2000)These regulators induce expression of CK response genes, including type-A *ARR*s, which in turn dampen the CK response and serve as indicators for CK pathway activation (To et al., 2004, 2008). In addition, several genes encoding enzymes involved in CK biosynthesis and degradation are among the CK response genes, resulting in complex, tissue specific feed-back systems.

While continuously active stem cells only contribute to postembryonic development of plants, apical meristems are set up and defined during embryogenesis, but remain inactive until germination. We have shown previously that during stem cell activation, *WUS* expression is synergistically controlled by photoreceptor-mediated light signaling pathways and photosynthesis derived sugars and that both pathways require activity of the TARGET OF RAPAMYCIN (TOR) kinase (Pfeiffer et al., 2016). TOR is an evolutionary conserved regulator of growth and acts as central nutrient sensor promoting anabolic processes, like protein biosynthesis under favorable conditions, while at the same time limiting autophagy driven catabolic turnover (Liu & Sabatini, 2020; Shi et al., 2018; Tafur et al., 2020). By phosphorylation of central regulators of ribosome biogenesis and cell cycle regulation, such as S6K1 and E2FA, TOR controls transcriptome and metabolic reprogramming to facilitate adaptation to changes in energy- and nutrient availability (Y. Dong et al., 2017; Xiong et al., 2013). Although much progress has been made in elucidating the TOR signaling landscape in mammals, our understanding of the TOR network in plants remains relatively poor. Recently, it has been demonstrated that TOR dependent phosphorylation of ETHYLENE INSENSITVE 2 (EIN2) controls growth of etiolated seedlings and mediates the transcriptional response to glucose in an ethylene independent manner (Fu et al., 2021). Moreover, the phytohormones auxin and abscisic acid have been identified as upstream regulators that influence TOR activity by modulating complex assembly with the associated regulatory protein RAPTOR (Li et al., 2017; Wang et al., 2018).

While the mechanisms connecting TOR with sugar and hormone responses have begun to emerge, little is known about how TOR controls shoot growth and SAM activity. We have shown previously that TOR dependent stem cell activation correlated with increased CK signaling in the SAM and that mutations in *ckx5* and *ckx6* promoted *WUS* expression during germination. However, the mechanisms connecting TOR kinase and CK signaling had remained elusive. Here, we report that for *WUS* expression and shoot development, the cytokinin species *tZ* is the most relevant downstream effector of TOR and demonstrate that translational regulation of cytokinin catabolic enzymes represents a major mechanism for TOR kinase to control levels of *tZ*.

## Results

### The transcriptional response to TOR inhibition in *Arabidopsis* shoots

Building on our previous results that demonstrated that TOR acts as the central integrator of light and sugar signaling during stem cell activation (Pfeiffer et al., 2016), we set out to identify downstream regulatory pathways that connect TOR with *WUS* expression and shoot development. Therefore, we analyzed the transcriptomes of *Arabidopsis* shoots with impaired TOR function. Because *tor null* mutants are lethal (Menand et al., 2002) and *Arabidopsis* is mostly insensitive to Rapamycin, we applied short term treatments with three independent TOR active site inhibitors, namely AZD8055, TORIN1 and KU63794 (Q. Liu et al., 2012; Montané & Menand, 2019; Schenone et al., 2011). We chose working concentrations (2 *µ*M AZD8055, 10 *µ*M TORIN1, 10 *µ*M KU63794) that have been shown to substantially impair shoot growth (P. Dong et al., 2015) and determined the earliest time point these inhibitors showed a robust effect on TOR activity in seedling shoots using S6K1 phosphorylation as a readout (Fig. S1a). We observed robust reduction of S6K1 phosphorylation in shoots of four days old seedlings eight hours after transfer to 2 *µ*M AZD8055 and 10 *µ*M TORIN1 but not for 10*µ*M KU63794. We therefore tested KU63794 at 20 *µ*M, which indeed led to substantially reduced S6K1 phosphorylation after 8h (Fig. S1b). Using these experimental parameters for our RNA-seq experiments, we were able to identify the expression of 23654 genes across all samples. Using DESeq2 (Love et al., 2014), we found 6639 differentially expressed genes (DEGs) in samples treated with a single inhibitor compared to their mock control (Fig. 1a + Table S1), among which 3266 (49.2%) showed reduced expression, and 3373 (50.8%) accumulated to higher levels than in controls. Most genes were affected by AZD8055 (5303 DEGs) followed by KU63794 (4556 DEGs) and TORIN1 (3692 DEGs). The different treatments caused significantly converging effects, with 2509 (37.7%) DEGs common to all three inhibitors and 4403 (66.3%) DEGs found in at least two of three treatments, suggesting that we were able to identify a set of robust TOR response transcripts. The differences between the inhibitors are likely due to unique selectivity-, potency and efficacy profiles of the three substances (Q. Liu et al., 2012; Montané & Menand, 2019; Schenone et al., 2011). Comparisons with previously published RNAseq data revealed significant overlap with studies using AZD8055 (P. Dong et al., 2015) or TORIN2 (Scarpin et al., 2020) as inhibitors (Figure S2), even though their experimental setups were substantially different. The overlap in DEGs compared with microarray data of RNAi lines against TOR three and six days after induction was rather mild (Caldana et al., 2013), which was expected given the highly divergent experimental setup.

**Figure 1:**
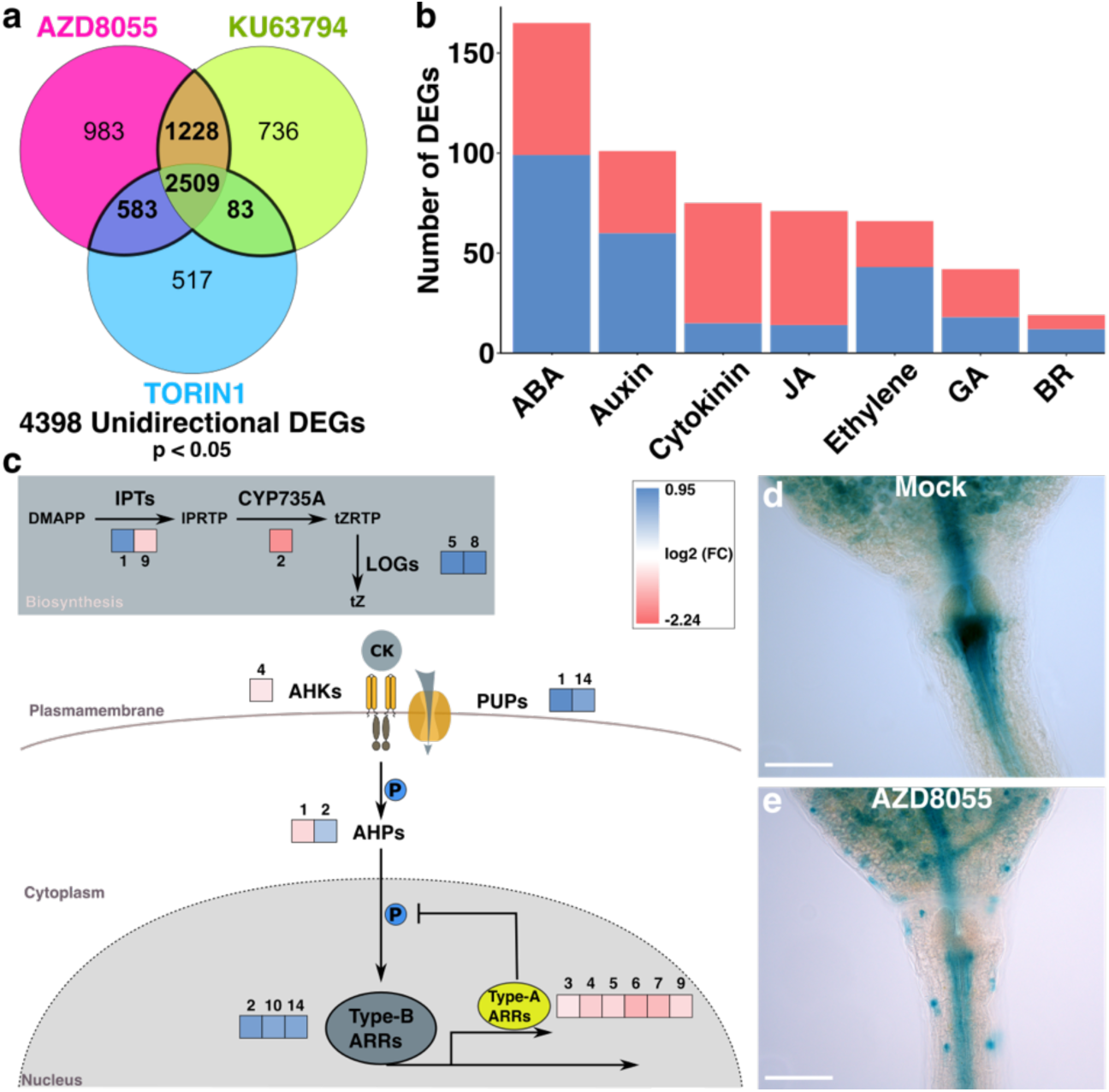
TOR inhibition leads to reduced CK signaling in the SAM. **a)** Venn diagram of differentially expressed genes obtained from RNAseq of shoot tissue from 4 day old seedlings treated with different TOR inhibitors for 8 h. **b)** Gene count of DEGs from RNAseq analysis annotated with hormone signaling function from GO term analysis. ABA = abscisic acid, JA = jasmonic acid, GA = gibberellic acid, BR = brassinosteroids. **c)** Schematic representation of the CK signaling pathway. Color code represents log fold change value obtained from RNAseq analysis. The numbers over the boxes indicate the isoform number of the respective gene. IPT = isopentenyltransferase, LOG = lonely guy, AHK = Arabidopsis histidine kinase, AHP = Arabidopsis histidine phosphotransferase, ARR = Arabidopsis response regulator, PUP = purine permease. **d, e)** Representative microscopic image of pTCSn:GUS reporter line treated with either DMSO or AZD8055 for 24 h. Scale bar = 30 *µ*m.

For further analysis we focused on mRNAs that showed a similar behavior in response to at least two inhibitors (significant difference in abundance with the same direction of change), which resulted in 4398 high confidence TOR target genes (Fig. 1a + Table S1). We chose to not apply any expression cut off because the genes selected were consistently changed in at least 6-9 replicates, rendering them interesting targets for investigation even if the fold change may have been low. Among the genes with reduced expression, GO categories (Table S1) of well characterized TOR dependent processes were most prevalent, such as translation, ribosome biogenesis, tRNA metabolism and anabolic processes (Dobrenel et al., 2016; Shi et al., 2018). The same applied to transcripts with increased accumulation among which we found GO categories (Table S1) related to catabolism, autophagy, secondary metabolism, glucosinolate biosynthesis and photosynthesis to be enriched (P. Dong et al., 2015; Malinovsky et al., 2017; Pu et al., 2017; Ren, 2015; Scarpin et al., 2020).

During detailed inspection of our data, we identified many DEGs with annotations related to hormone signaling pathways (Fig. 1b + Table S1) with genes related to abscisic acid (ABA) but also auxin, cytokinins (CKs), jasmonic acid (JA), ethylene, brassinosteroids (BR) and gibberellic acid (GA) being well represented. Interestingly, for most hormones the number of transcripts showing increased or reduced levels were similar. Notable exceptions were CKs and jasmonic acid, where the abundance of most transcripts was reduced in response to TOR inhibition. This apparent reduction of pathway activity caught our attention because of the known positive effect of CKs on SAM activity and shoot development (Kiba et al., 2013; Osugi et al., 2017). The transcriptome analysis revealed differential regulation of genes involved in CK biosynthesis (*IPT1, −9. CYP735A2, LOG5, −8*), -transport (*PUP1, −14*), -sensing (*AHK4*), - signal transmission (*AHP1, −2*) and -transcriptional regulation (*ARR2, −10, −14*) (Fig. 1c). Notably, all six type-A *ARRs* that appeared as differentially regulated in our dataset were repressed after inhibitor treatment. Expression of type-A *ARRs* is commonly used as an approximation for CK signaling output and we thus concluded that CK signaling response is likely reduced following TOR inhibition.

### TOR inhibition reduces cytokinin signaling in the SAM

To test whether our findings obtained at the level of the entire seedling shoot bear any relevance for the SAM, we used the well-established *pTCSn:GUS* CK signaling reporter line. Supporting the RNA expression data for type-A *ARRs*, we found that *pTCSn* activity was substantially reduced after AZD8055 treatment especially in the meristematic region of the shoot, where CK function is required to maintain *WUS* expression (Fig. 1d+e + Fig. S3). We next asked whether the reduction in CK signaling would also affect *WUS* expression. To this end, we utilized our previously established *pWUS:3xVenus:NLS* transcriptional reporter system (Pfeiffer et al., 2016). Importantly, we investigated etiolated seedlings, since in their SAMs *WUS* is usually not expressed giving us a maximum of sensitivity to test different stimuli for their ability to activate *WUS*. In addition, we have previously shown that the CK 6-benzyladenine (6-BA) is able to induce *WUS* expression in this setting (Pfeiffer et al., 2016), allowing us to quantitatively test the effect of TOR inhibition on CK dependent activation of *WUS*. Indeed, CK dependent activation of *WUS* was almost fully blocked when 6-BA was applied together with AZD8055 (Fig. 2a). Consistently, 6-BA induced expression of several type-A *ARRs* was suppressed when seedlings were preincubated with the TOR inhibitor (Fig. S4). Together, this suggested that TOR inhibition interferes with the activation of various CK response genes in the SAM, including *WUS*.

**Figure 2:**
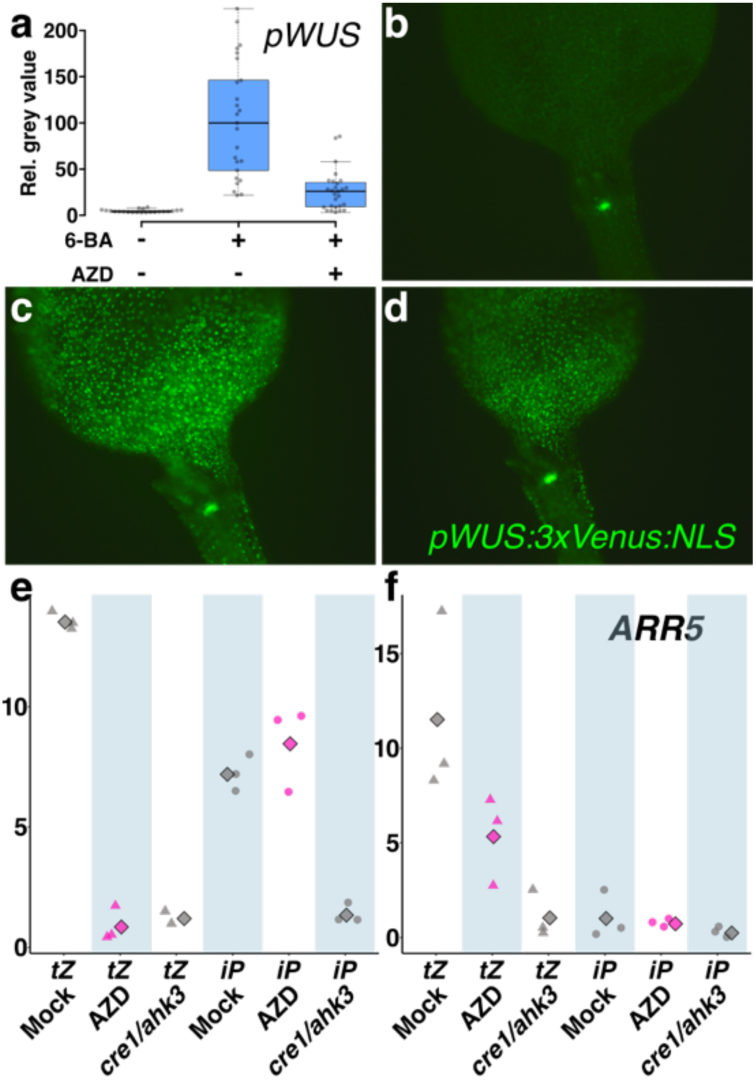
CK output potential is not affected by TOR inhibition. **a)** Quantification of pWUS:3xVenus:NLS reporter signal from 5 day old etiolated seedlings treated with 0.5 *µ*M 6-benzyladenine and 2 *µ*M AZD8055 for 3 days. Significant differences between Mock-BA(p=4.05e-13), Mock-BA+AZD (p=5.88e-08) and BA-BA+AZD (p=1.42e-07) were calculated with Wilcoxon rank sum test with Hochberg correction. n=22-27 **b-d)** Representative microscopic images of p35S:ARR1∂DDK:GR crossed with pWUS:3xVenusNLS. 3 day old light grown seedlings were treated with b) mock, c) 10 *µ*M dexamethasone (DEX) or d) DEX + 2 *µ*M AZD8055 for 24 h. **e, f)** Relative expression values, normalized to PP2A obtained with q-RT-PCR from e) root and the respective f) shoot tissue are shown. 4 day old seedlings preincubated on AZD8055 or mock for 8h and subsequently sprayed with either 100 nM trans-zeatin (*t*Z) or 100 nM isopentyladenine (*i*P) solution for 30 min. Data points show expression values from biological replicates (triangles and circles) together with the calculated mean (rhombus).

Since reduced CK signaling could be a major cause for the previously observed effects of TOR inhibition on SAM activity and *WUS* expression, we next aimed at identifying at which step of the pathway this effect may occur. Transcription factors of the type-B ARRs family have been shown to directly bind to the *WUS* promoter and induce *WUS* expression in response to CK sensing (Dai et al., 2017; Gruel et al., 2016; Meng et al., 2017). To test the functional interaction between TOR and type-B ARRs for *WUS* activation, we crossed our *pWUS:3xVenus:NLS* reporter with a line carrying an inducible version of a constitutively active allele of ARR1 (*p35S:ARR1ΔDDK:GR*). This version was expressed from a strong and ubiquitous viral promoter (*p35S*) and the transcription factor was lacking the receiver domain (*ΔDDK)*, making it independent of CK induced phosphorylation. In addition, it was fused to a glucocorticoid receptor (GR) domain, enabling dexamethasone (DEX) dependent nuclear translocation and thus experimental induction (Sakai et al., 2001). Upon DEX treatment of *p35S:ARR1ΔDDK:GR* shoots, *WUS* became expressed in most cells (Fig. 2b+c+d + Fig. S5) confirming that it is a direct type-B ARR target gene. Interestingly, treatment with AZD8055 did not interfere with this response, indicating that TOR interacts with CK signaling upstream of type-B ARRs. Moreover, we found that ARR1ΔDDK mediated induction of type-A *ARR* expression was quantitively not affected by TOR inhibition (Fig. S6), which further supported that type-B ARR transcriptional activity is independent of TOR.

To further narrow down the intersection of TOR kinase activity and CK signaling, we assessed whether seedlings would still be able to respond to treatment with endogenous CKs after TOR inhibition. We therefore performed CK response assays, using *tZ* and *iP*, mRNA levels of *ARR5* as readout, and the CK receptor double mutant *cre1-2/ ahk3-7,* which has impaired sensitivity to *tZ* and *iP*, as negative control (Riefler et al., 2006) (Fig. 2e+f). Moreover, we decided to include roots in our assay system since *iP* and *tZ* are known to evoke different responses in shoots and roots. To resolve even subtle differences in CK sensing we decided to use relatively low CK concentrations with 100 nM *tZ* and *iP*.

Despite the low concentration, *tZ* application evoked a strong increase of *ARR5* expression compared to the *cre1-2/ ahk3-7* mutant indicating a solid hormone response (Fig. 2e+f). However, when the seedlings were pre-incubated with AZD8055, no difference in *ARR5* expression compared with the *cre1-2/ ahk3-7* mutant was observed in roots and reduced response in shoots (Fig. 2e+f), suggesting that TOR activity is required for transcriptional responses to exogenously applied *tZ*. Interestingly, treatment with the CK derivative *iP* had different effects. Whereas shoots seemed to be generally insensitive to this low concentration of *iP* (Fig. 2f), wt roots showed solid *ARR5* accumulation compared to the *cre1-2/ ahk3-7* mutant (Fig. 2e). Notably, treatment with AZD8055 did not interfere with this response, suggesting that seedlings are in principle still able to sense CKs after TOR inhibition. Since CRE1 and AHK3 CK receptors have higher affinities towards *tZ* than *iP* (Romanov et al., 2006), it appeared unlikely that the differential response observed after TOR inhibition could arise at the receptor or the signal transduction level.

### Reducing TOR activity affects cytokinin homeostasis

Based on the differential CK responses described above, we hypothesized that the functional intersection of TOR and CK must be affecting substrate availability, by either selective degradation or sequestration of specific CKs, such as *tZ*. We therefore used metabolite analyses to quantify the abundance of CK molecules in response to TOR inhibition and found drastic changes in CK homeostasis following AZD8055 treatment (Fig. 3). The results revealed that total *t*Z levels were significantly reduced, whereas *i*P metabolites were not affected. This confirms the tZ specific effects of AZD8055 from our CK response assay (Fig. 2e+f). On the level of individual CK metabolites *iP* active bases were enriched two-fold, whereas the monophosphate precursor (*iP*RMP) was slightly depleted. The changes in *tZ* derivatives were even more pronounced, and we observed a reduction of active *tZ* bases to 50% of the mock levels after AZD8055 treatment. Moreover, a four-fold decrease of *trans-*zeatin riboside (*tZ*R) and a six-fold decrease of the monophosphate precursor (*tZ*RMP) occurred in response to TOR inhibition, whereas the conjugated glycosides *tZ*OG and *tZ*9G were almost unchanged for both *tZ* and *iP* derivatives. Only tZ7G also showed a significant reduction of about 30%. In several studies *tZ* and *tZ*R have been shown to be the main CKs driving shoot development and *WUS* expression (Kasahara et al., 2004; Kiba et al., 2013; Landrein et al., 2018; Osugi et al., 2017) and thus this drastic reduction in *tZ* and *tZ*R upon TOR inhibition fitted well with effects of TOR activity on *WUS*. Strikingly, *cis*-zeatin (*c*Z) active bases accumulated six-fold as well as all other *c*Z derivatives (Fig. 3). However, *c*Z does not show relevant CK activity in classical bioassays (Gajdošová et al., 2011; Kasahara et al., 2004) and in line with this it had very little effect on our *WUS* reporter in etiolated seedlings compared with *tZ* and *iP* (Fig. S7).

**Figure 3:**
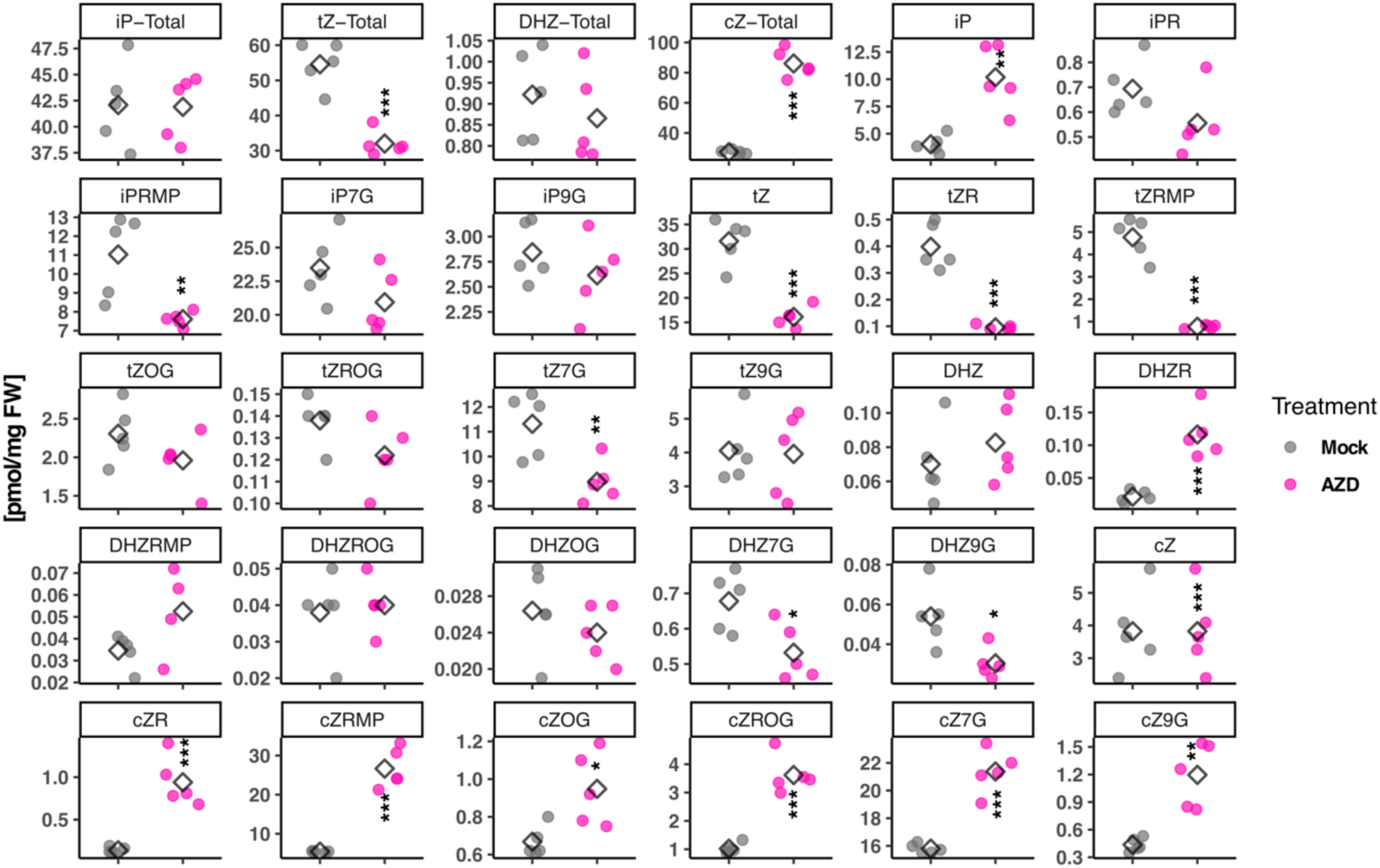
Disturbed CK homeostasis after TOR inhibition. Quantification of different CK compounds in 4 day old seedlings treated with AZD8055 (magenta) or mock (grey) for 8 h. Asterisks indicated significant differences of the AZD8055 treated sample compared with the respective mock calculated with two-tailed t-test (*, **, and *** correspond to p-values of 0.05 > p > 0.01, 0.01 > p > 0.001, and p < 0.001, respectively). Error bars indicate standard deviation. n =5. **a)** *t*Z = trans-zeatin, *t*ZR = trans-zeatin riboside, *t*ZRMP = trans-zeatin riboside-5’monophosphate, *t*ZOG = trans-zeatin O-glucoside, *t*ZROG = trans-zeatin riboside O-glucoside, *t*Z7G = trans-zeatin-7-glucoside, *t*Z9G = trans-zeatin-9-glucoside **b)** *i*P = isopentenyladenine, *i*PR = isopentenyladenosine, *i*PRMP = isopentyladenosine-5’monophosphate, *i*P7G = isopentyladenine-7-glucoside, *i*P9G = isopentyladenine-9-glucoside **c)** DHZ = dihydrozeatin, DHZR = dihydrozeatin riboside, DHZRMP = dihydrozeatin riboside-5’monophosphate, DHZOG = dihydrozeatin O-glucoside, DHZROG = dihydrozeatin riboside O-glucoside, DHZ7G = dihydrozeatin-7-glucoside, DHZ9G = dihydrozeatin-9-glucoside **d)** cZ = cis-zeatin, cZR = cis-zeatin riboside, cZRMP = cis-zeatin riboside-5’monophosphate, cZOG = cis-zeatin O-glucoside, cZROG = cis-zeatin riboside O-glucoside, cZ7G = cis-zeatin-7-glucoside, cZ9G = cis-zeatin-9-glucoside

### Cytokinin and sugars act downstream of TOR

Based on the striking differential accumulation of *tZ* and *i*P after TOR inhibition and the divergent potential of these molecules to cause CK responses in the presence of AZD8055, we hypothesized that TOR controls shoot development via the regulation of *tZ* levels. We reasoned that if reduced *tZ* availability was causal for the reduced *WUS* expression observed under TOR inhibition, exogenous re-supplementation should be able to rescue *WUS* promoter activity. Indeed, when we supplemented AZD8055 treated seedlings with 0.5 *µ*M of different CKs, we observed that only *tZ*R, the synthetic CK 6-BA and to a lesser extent also *tZ* were able to rescue *WUS* expression under TOR inhibition (Fig. 4a-e). This was in agreement with previous studies, showing that *tZ*R and 6-BA have stronger potential to induce *WUS* and shoot development compared to *tZ* (Landrein et al., 2018; Osugi et al., 2017). However, at higher concentrations also *iP* reverted *WUS* expression to wt levels (Fig. S8).

**Figure 4:**
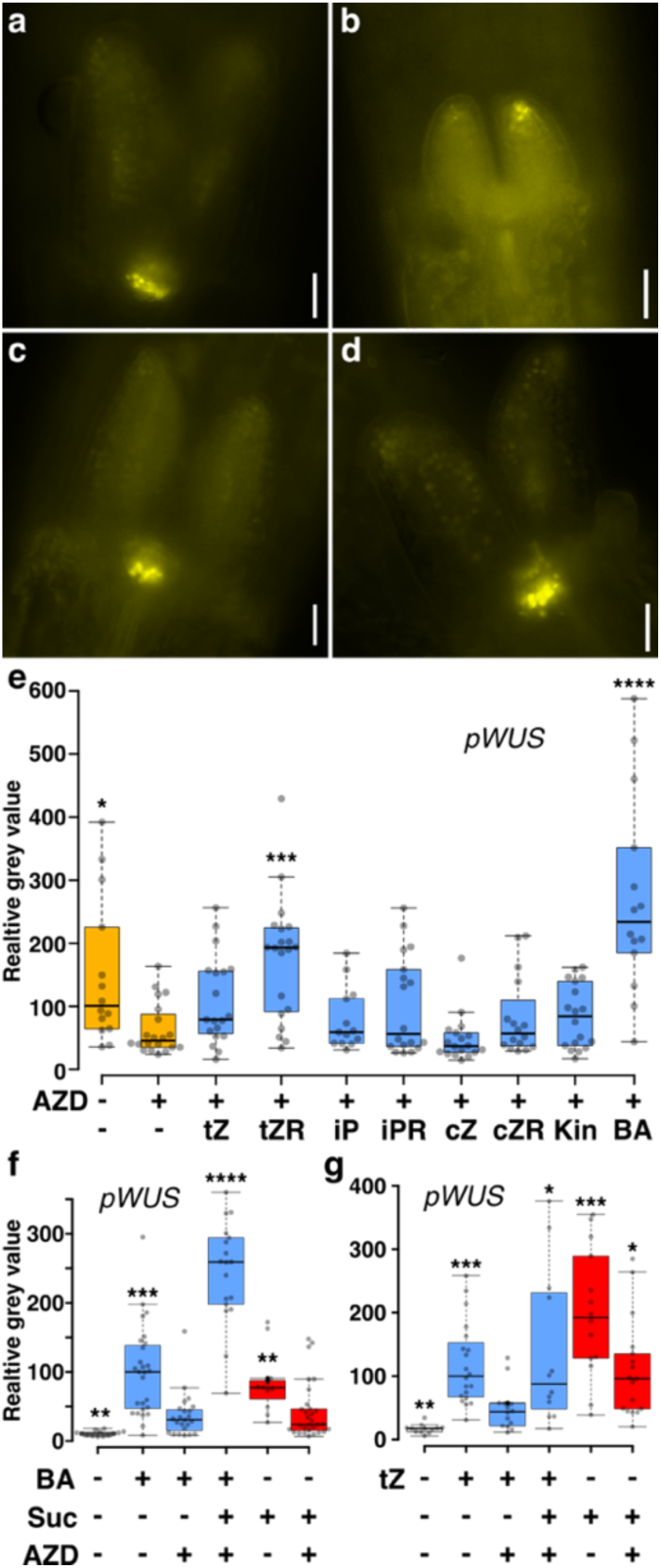
trans-zeatin derivatives rescue *WUS* expression upon TOR inhibition. Representative microscopic images of e) pWUS:3xVenus:NLS reporter line treated with **a)** mock **b)** 2*µ* AZD8055 **c)** AZD + 0.5 *µ*M *t*ZR and **e)** AZD + 0.5 *µ*M 6-BA. Images were acquired as black and white and false color coded with imagej. Scale bar = 30 *µ*m. **e)** Quantification from a)-d) of pWUS:3xVenus:NLS reporter signal. 3 day old seedlings grown in the light were treated with 2 *µ*M AZD8055 and 0.5 *µ*M of different CK derivatives for 1 day. Asterisks indicate significant differences compared to the AZD treated conditions. n=13-20. **f, g)** Quantification of pWUS:3xVenus:NLS reporter signal. 2 day old etiolated seedlings were treated with 0.5 *µ*M 6-benzyladenine (BA) or trans-zeatin (*t*Z), 2 *µ*M AZD8055 and/or 1% sucrose for 3 days in the dark. Asterisks indicate significant differences compared to the BA+AZD or *t*Z+AZD condition. **f)** n=13-30 **g)** n=12-20. Significance levels were calculated using the Wilcoxon rank sum test with Hochberg correction (* = p < 0.05; ** = p < 0.01;*** = p < 0.001, **** = p < 0.0001). *t*Z = trans-zeatin, *t*ZR = trans-zeatin riboside, *i*P = isopentenyladenine, *i*PR = isopentenyladenosine, cZ = cis-zeatin, cZR = cis-zeatin riboside, Kin = kinetin, BA = 6-benzyladenine.

Interestingly, this behavior was different to the one observed during stem cell activation in dark grown seedings, where 6-BA induced *WUS* expression was suppressed when TOR is inhibited (Fig. 2a). We therefore hypothesized that reduced availability of photoassimilates might be responsible for the different results obtained with etiolated seedlings. Indeed, when we supplied sugar together with 6-BA or *tZ* under TOR inhibitory conditions in dark grown seedlings, *WUS* expression was elevated to levels even higher than with 6-BA alone (Fig. 4f+g). This result was striking, since it implied that while TOR is the central gatekeeper for light-, sugar- or CK dependent activation of *WUS*, respectively, once sugars and CK act together, they are sufficient to drive *WUS* expression downstream of TOR activity. Noteworthy, the effect of sucrose alone varied between the experiments, but was not significantly different compared to *tZ*+AZD+Sucrose (p = 0.2, Students t-test). In sum, our results so far demonstrated that TOR inhibition leads to strongly reduced *tZ* content in shoots, which in turn is causal for reduced expression of *WUS*.

Since the relationship between TOR and *WUS* appeared to be mainly governed by *tZ* availability, we next wanted to identify the mechanisms underlying differential *tZ* accumulation. The observed reduction of *tZ* derivatives could be the result of either reduced biosynthesis or increased turnover and our data so far supported both mechanisms. On the one hand, expression of the gene encoding the *tZ* synthesis enzyme *CYP735A2* (Takei et al., 2004) was substantially reduced following AZD8055 treatment (Fig. 1c, Table S1), suggesting biosynthesis may be causal. However, transcriptional repression of *CYP735A2* does not allow any conclusion on the protein levels or enzymatic activity and moreover a second isoform CYP735A1 exists which has been shown to compensate for the loss of the other isoform (Kiba et al., 2013). On the other hand, shoots showed impaired response to exogenous *tZ* treatment upon TOR inhibition (Fig. 2e+f), clearly pointing towards increased *tZ* turnover after TOR inhibition. Importantly, inhibition of the biosynthesis pathway at various steps should lead to clearly identifiable footprints in the accumulation levels of the diverse CK species. However, the increase of *iP* after inhibition was in the range of 5 pmol/mg FW whereas *tZ* reduction was about 15 pmol/mg FW, suggesting that a shift between this species is unlikely. In addition, conversion from *iP* to *tZ* happens mostly on the precursor levels as CYP735A highly favors *iP*RMP as a substrate and has almost no affinity towards *iP* (Takei et al., 2004) and *iP*RMP levels decreased following AZD8055 treatment (Fig. 3b) further pointing towards increased turnover. Another explanation could be interconversion from *tZ* to *c*Z by *cis-trans* isomerase, as *c*Z is drastically increased after TOR inhibition. However, the increase of all *c*Z compounds is much higher compared with the decrease of the respective *tZ* compounds. More importantly, the existence of such an enzyme in plants has not been shown and appears highly unlikely (Hluska et al., 2017; Miyawaki et al., 2006).

### TOR activity controls cytokinin turnover via CKX enzymes

To elucidate the relative contribution of reduced biosynthesis and increased turnover, we decided to first investigate the role of CKX cytokinin degrading enzymes in TOR mediated control of shoot development and regulation of *WUS.* To this end, we employed shoot fresh weight analyses as a measure of growth in response to treatment with increasing concentrations of AZD8055. These simple assays allowed us to determine effective doses for growth inhibition (ED) across a range of genotypes and revealed that CKX enzymes are at least partially responsible for the observed effect of TOR on cytokinin mediated growth control (Fig. 5a+b+c+e, Fig.S9-S12). First inspection of the dose response curves suggested that wild type seedlings were more sensitive to low concentrations of AZD8055 compared with the *ckx* mutant seedlings, whereas in the higher concentration range the curves converge (Fig. 5a+b+c+e). Statistical analysis revealed that the wt shoot fresh weight differed significantly between 0 and 0.5 *µ*M AZD8055 and between 0.1 and 0.5 *µ*M AZD indicating that the growth inhibitory effect for wt seedlings kicked in between 0.1 and 0.5 *µ*M. In contrast, shoot fresh weight for the *ckx1, ckx2*, *ckx3*, *ckx4*, *ckx5*, *ckx6, ckx1/3* and *ckx1/3/5* mutants did not differ significantly between 0 and 0.5 *µ*M but was different between 0.5 and 1 *µ*M, suggesting that growth inhibition for these mutants only occurred above 0.5 *µ*M AZD8055. Noteworthy, *ckx1* showed resistance up to 1 *µ*M AZD8055. The *ckx5/6* mutant appeared to behave largely like wt seedlings.

**Figure 5:**
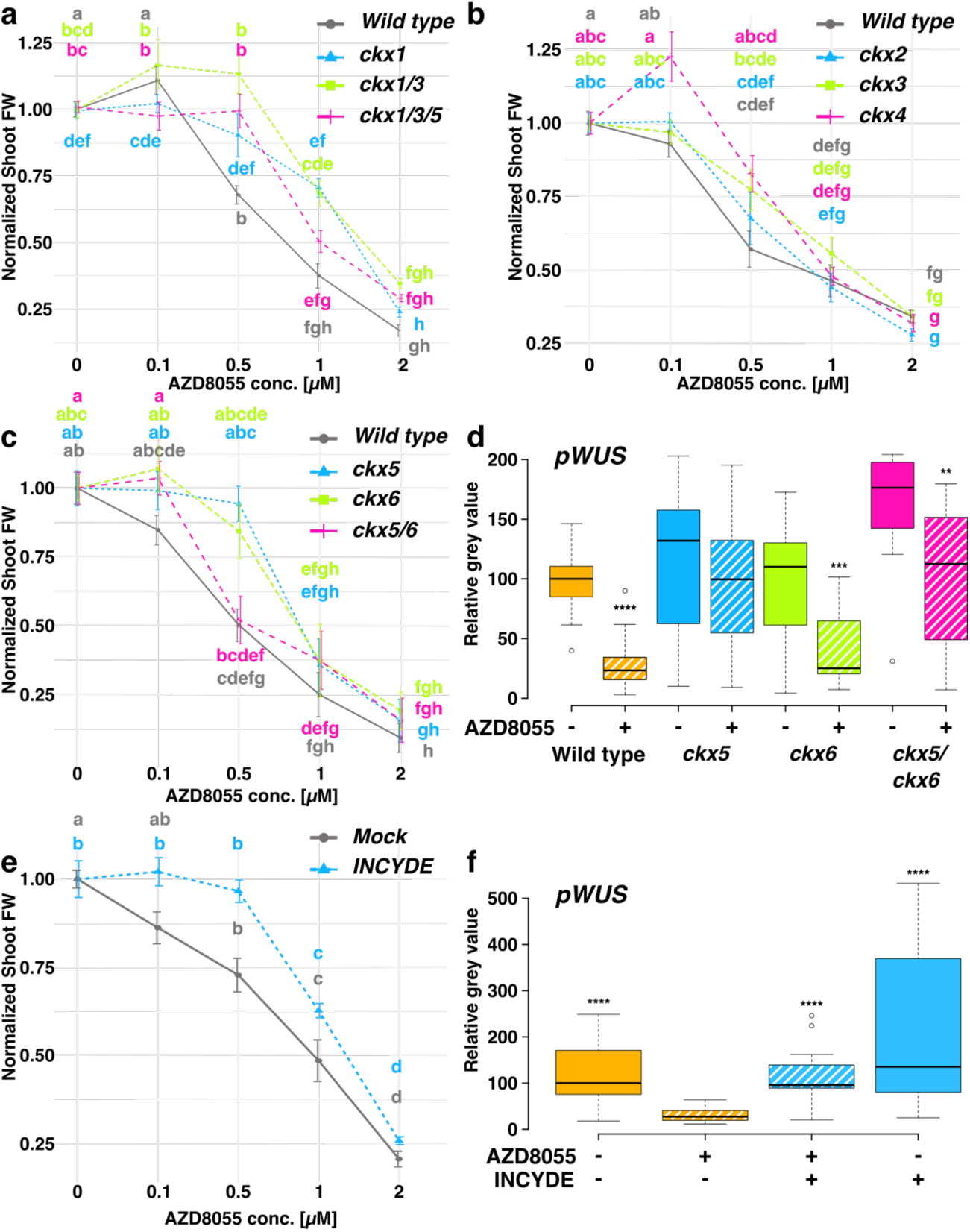
Specific *CKX* mutants are resistant to TOR inhibition. **a, b, c, e)** Average shoot fresh weight per seedling was calculated from measurements of 40 seedlings in batches of 5-10 for each measurement, that were transferred at 4 DAG to different concentrations of AZD8055 for 7 days. Data was pooled from two independent experiments and normalized to untreated mean. Error bars represent standard error. Unnormalized data is found in the supplements. Significant interaction was tested before normalization with ANOVA linear models and post hoc t-test with bonferoni correction. Letters indicate significance levels **c)** *ckx5*, *ckx6* and *ckx5/6* are in the genetic background of *pWUS:3xVenusNLS*, *pCLV3:mCherry:NLS* and were tested against this background. **e)** 75 nM INCYDE. **d, f)** Quantification of pWUS:3xVenus:NLS reporter signal. Seedlings were grown in the light for 3 days and treated with mock, 2 *µ*M AZD8055 and/ or 75 nM INCYDE for 1 day. **d**) Asterisks indicate significant differences between the AZD treated condition and the respective Mock. n=15-29 **f)** Asterisks indicate significant differences compared to the AZD treated condition. n=16-19. For d) and f) Significance levels were calculated using the Wilcoxon rank sum test with Hochberg correction.

These observations were corroborated by inspection of the different ED50 ratios (Fig. S12). ED50 describes the effective dose of AZD8055 to achieve 50% growth inhibition and the ratios show that all *ckx* mutants had a higher ED50 compared to wt. However, for *ckx2* and *ckx5/6* mutants the confidence interval overlapped with the wt, suggesting that those two mutants exhibited a wt like response.

Taken together, these experiments suggested that multiple CKX enzymes were likely to contribute to the effect of TOR on CK signaling. Importantly, this notion was also supported by experiments employing the CKX inhibitor INCYDE (Zatloukal et al., 2008). Treatment with 75 nM INCYDE made wild type seedlings fully resistant to even 0.5*µ*M AZD8055 (Fig. 5e), demonstrating that the enhanced activity of multiple CKX enzymes was responsible for mediating the control of shoot growth by the TOR kinase. However, our results also demonstrated that reduced CKX activity did not render seedlings resistant to full block of TOR kinase activity, which may be a physiologically rare state. Moreover, it was evident that there are specific contributions of individual CKX isoforms which was highlighted by the loss of AZD8055 resistance in the *ckx5/6* double mutant compared to the respective single mutants, which pointed towards a complex genetic interaction.

To connect shoot growth with SAM activity and to investigate the role of *CKX* genes in controlling *WUS* expression in response to TOR, we used CRISPR alleles of *CKX5* and *CKX6* we had generated previously in the *WUS* reporter background, because *WUS* RNA levels in seedlings cannot be quantified reliably by other means. Supporting our findings at the level of shoot growth, we found that inactivation of *CKX5*, or *CKX5* together with *CKX6* completely restored *WUS* expression to wild type levels after TOR inhibition, whereas mutation of *CKX6* alone had no effect (Fig. 5d). The untreated *ckx5,6* double mutant exhibited higher basal expression levels of the *WUS* reporter which were reduced upon AZD8055 treatment but not below wild type mock levels, demonstrating additive genetic interaction between *ckx5* and *ckx6* independent of TOR input. Again, as for shoot growth, pharmacological inhibition of CKX activity using INCYDE (Zatloukal et al., 2008) also rescued *WUS* expression to mock levels in the presence of 2 *µ*M AZD8055 (Fig. 5f). Together, these observations demonstrated that increased CKX activity upon TOR inhibition caused reduced *WUS* expression and in turn impaired shoot growth, pointing towards a molecular interaction between TOR and the CKX enzyme family.

### TOR inhibition leads to CKX1 protein accumulation

TOR could affect CKX enzymes at various levels from expression of the corresponding genes to *CKX* mRNA translation, to enzyme activity, or CKX protein stability. Mining our transcriptome data, we found no evidence of differential regulation of *CKX* genes at the transcript level, which we were able to confirm by RT-qPCR (Fig. S13). Only *CKX6* showed a mild but significant increase in transcript abundance, whereas *CKX2* and *CKX5* transcripts were even decreased. Hence, we concluded that the nature of the molecular interaction between TOR and CKXs likely occurs downstream of the mRNA level. To distinguish between increased protein accumulation and CKX enzyme activity, we analyzed protein abundance in response to AZD8055. Since there are currently no antibodies against any of the CKX enzymes available, we made use of a *p35S:cMyc-CKX1* translational fusion line (Niemann et al., 2015, 2018). Western blot analysis revealed that eight hours after transfer to AZD8055 containing medium, shoots accumulated 39 – 109 % more cMyc-CKX1 than mock treated shoots in three independent experiments (Fig. 6a). In contrast, AZD8055 did not induce significant changes in *CKX1* transcript abundance in the investigated line (Fig. 6b) during the experiment. Protein accumulation is either the result of decreased protein turnover or increased protein biosynthesis and hence we first tested if CKX protein stability was affected by TOR inhibition. To this end, we utilized cycloheximide (CHX) to inhibit translational elongation and assayed cMyc-CKX1 protein levels over the course of eight hours with- or without preincubation on AZD8055 for eight hours (Fig. 6c). We found that CKX1 turnover rate was very similar in AZD8055 and mock treated shoots and the protein half-life was determined at 4 h for both, which is in agreement with published results (Niemann et al., 2015). As a result, we concluded that TOR inhibition does not affect CKX1 protein stability but may rather enhance CKX1 translation to cause the observed protein over-accumulation (Fig. 6a).

**Figure 6:**
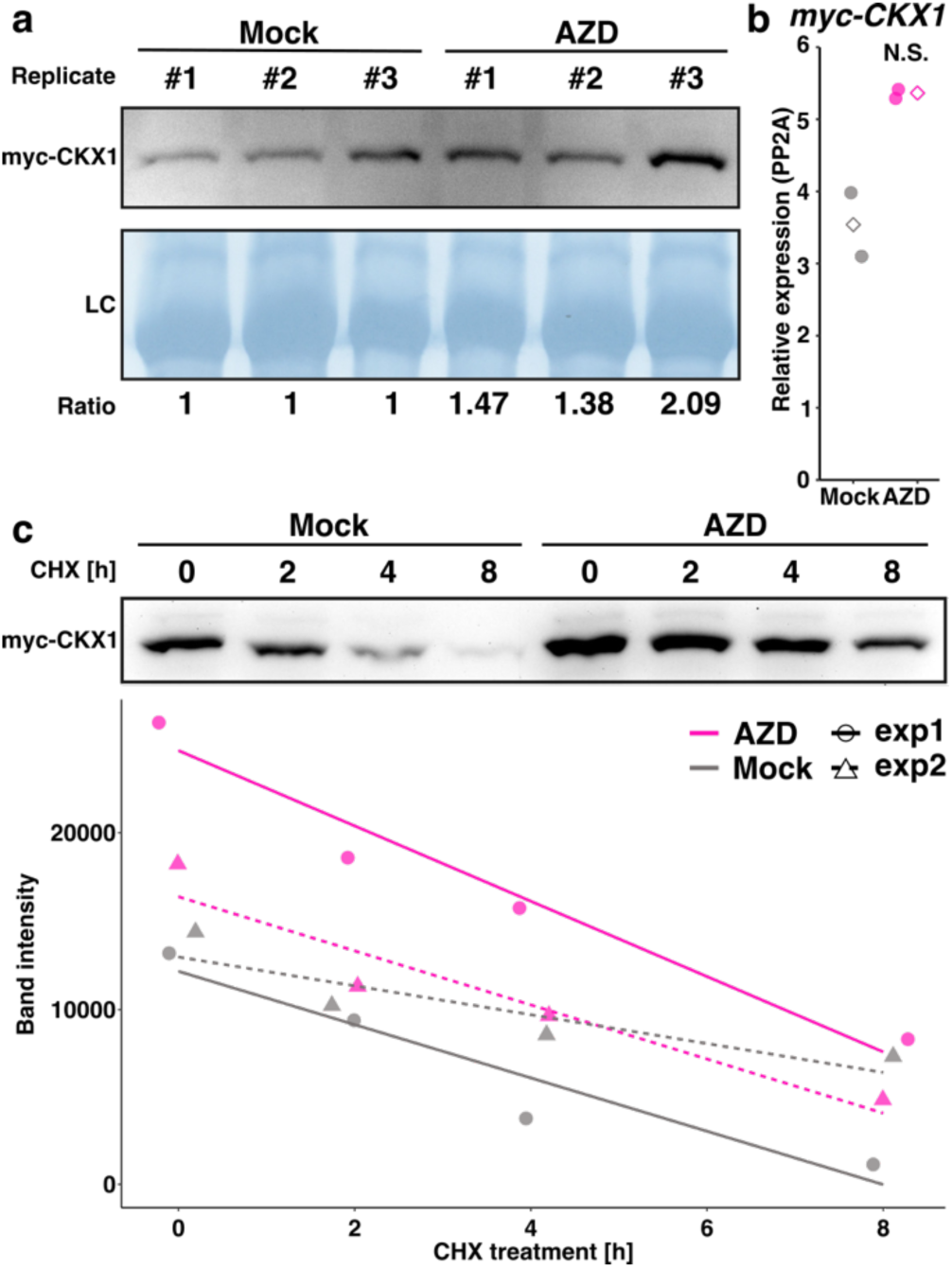
cMyc-CKX1 protein accumulates upon TOR inhibition. **a)** Western blot of p35S:cMyc-CKX1 4 day old seedlings treated with either mock or 2*µ*M AZD8055 for 8h. Shoot protein extract was probed with anti-cMyc serum. Replicates are from three independent experimental repetitions. Loading control (LC) stained with amido black. Ratios were calculated between band intensities from myc-CKX1 signal and the LC and normalized against the respective mock sample. **b)** q-RT-PCR of p35S:cMyc-CKX1 shoots with primers against the ectopic transcript. No significant difference using two biological replicates was found using paired t-test. Rhomboid indicates median value. **c)** Western blot of p35S:cMyc-CKX1 shoot protein extract probed with anti cMyc serum. Seedlings were pre-incubated on AZD8055 and then flooded with 200 *µ*M cycloheximide solution for the indicated time. Band intensities from two independent experimental repetitions were plotted with linear regression lines.

### TOR controls translation of specific *CKX* transcripts

To test this directly, we applied polysome profiling to analyze whether TOR inhibition modulates the association of different *CKX* mRNAs with highly translating polysomes (Fig. 7 + S14 + S15). To this end, we prepared polysome fractions from mock and AZD8055 treated shoots using sucrose gradient centrifugation and quantified the abundance of *CKX* mRNAs in the fractions using RT-qPCR. Consistent with the enhanced protein accumulation of CKX1 and AZD8055 resistance of *ckx1* mutants, we observed a strong enrichment of *CKX1* mRNA in the heavy polysome fractions (fractions 1-3), which suggested that more CKX1 protein was synthesized at the ribosomes after eight hours of TOR inhibition compared to mock treated controls. Similarly, we observed accumulation of *CKX5* and *CKX3* transcripts in polysomal fractions (Fig. 7), which fitted well with the observed resistance of the *ckx5* and *ckx3* mutants towards AZD8055 treatment (Fig. 5), although we were only able to detect *CKX3* transcript in one of three replicates tested. We also observed subtle accumulation of *CKX6* and *CKX7* transcripts in some of the lighter polysomal fractions (fractions 3-5) following AZD8055 treatment (Fig. 7 + S15). The lower enrichment of *CKX6* mRNAs in polysomes compared to *CKX5* fitted well with the behavior in our WUS reporter assays observed for the respective mutants (Fig. 5). In the case of *CKX4*, it was difficult to draw a clear conclusion as in some replicates it accumulated in polysomal fractions whereas in other replicates it was depleted from polysomes (Fig. 7 + Fig S15). In conclusion, our experiments showed that translation of multiple transcripts encoding CKX enzymes was controlled by TOR activity. In particular CKX1, CKX3 and CKX5 protein biosynthesis appeared to be repressed when TOR is active, which fitted well with functional data on shoot growth controlled by CKX1, CKX5 and CKX3 (Fig. 5) and with protein analysis of CKX1 (Fig. 6). In contrast, translation of *CKX4* appeared unaffected or even enhanced by TOR activity (Fig. 7 + Fig S14), whereas our genetic data showed a clear antagonistic effect of both players (Fig. 5 + Fig. S12). Close inspection of the 5’-leader sequences of all *CKX* encoding genes revealed that only *CKX4* contains a strong uORF signature that may influence the observed response to TOR inhibition at the translational level.

**Figure 7:**
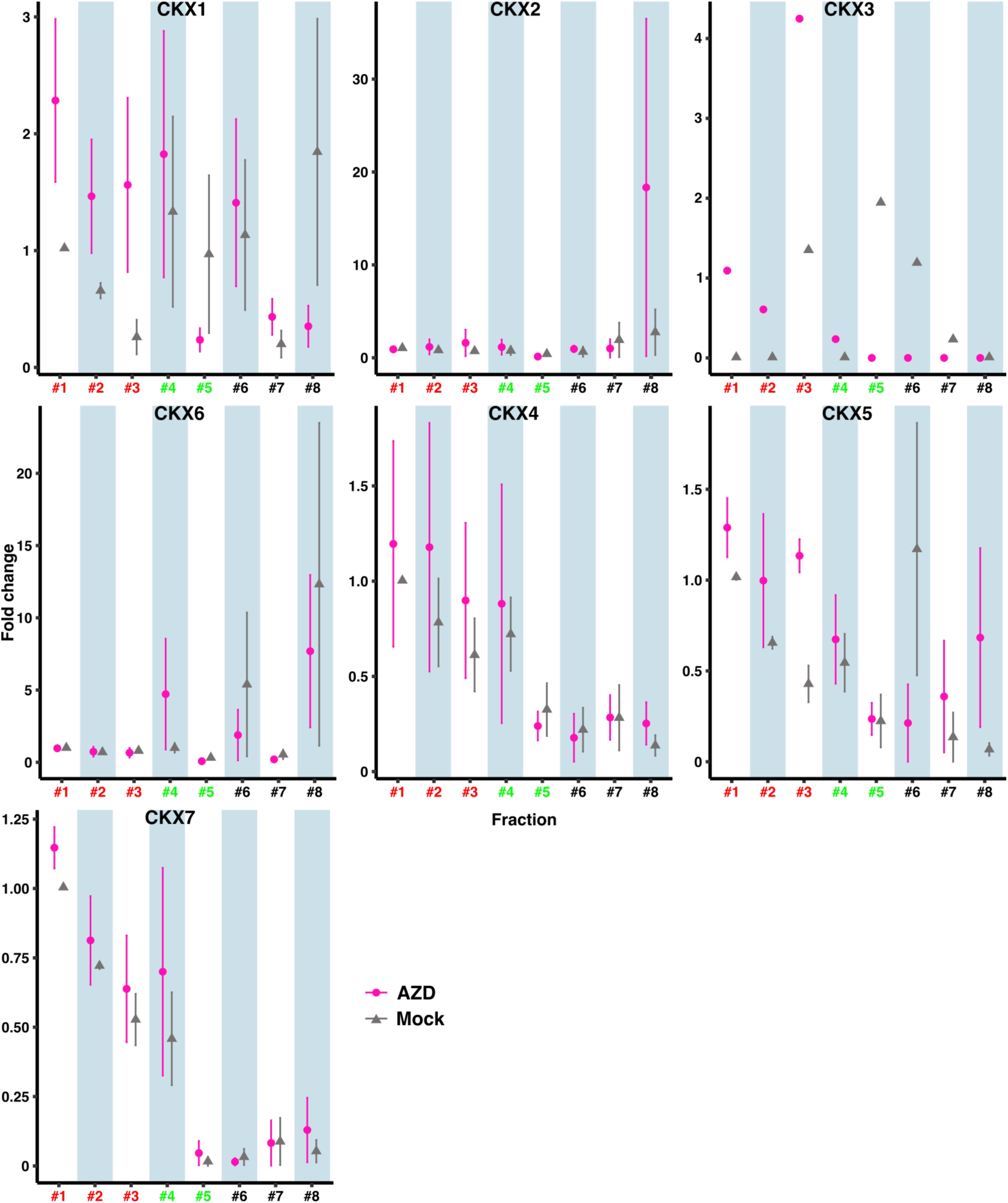
Transcripts of specific CKX isoforms accumulate in heavy polysomal fractions. CKX transcripts were quantified in ribosomal fractionations by q-RT-PCR relative to *UBI10* and normalized to the respective mock of fraction #1. Error bars represent standard error of the mean of data pooled from three independent experimental repetitions (except CKX2+7 were only detected in 2 replicates, CKX3 was detected in 1 replicate). Red numbers indicate heavy polysomal fractions (fractions 1-3), green numbers light polysomal fractions (fractions 4+5) and black number monosomal fractions (fractions 6-8). Individual data points are shown in the supplements (Fig. S18).

### *ckx1* mutants are fully resistant to TOR mediated reduction on *trans*-Zeatin metabolites

So far, our results clearly showed that CKXs play an important role mediating the effects of reduced TOR activity on shoot development. However, how *ckx* mutations affect CK metabolism in response to TOR inhibition and to which extent reduced *tZ* biosynthesis is involved remained unanswered. We therefore decided to evaluate the effect of AZD8055 on CK metabolites in *ckx1* and *ckx1/3* mutants (Fig. 8 + S16 + S17). Inspection of total levels of *i*P and *t*Z after AZD8055 treatment confirmed the previously observed reduction of *t*Z and the absence of an effect on *i*P levels (Fig. 8), which was consistent with our qRT results showing a specific effect on *t*Z but not *i*P (Fig. 2e+f). Strikingly, the total levels of *tZ* metabolites in *ckx1* mutants remained unchanged after TOR inhibition. This clearly shows that at the investigated time point, catabolic turnover of *t*Z by CKX1 is the dominant mechanism mediating the effects of TOR inhibition. The *ckx1/3* double mutant had initially higher *t*Z levels compared to the wt, which were reduced upon AZD8055 treatment, however, not below wt mock levels. In contrast, neither the total *i*P level, nor its response was affected in the two mutants. Inspecting the levels of individual CK metabolites, it became evident that the major effect of *ckx* mutations on *t*Z was on its N- and O-glucosides (Fig. S16). Particularly, the highly abundant *t*Z7G and *t*Z9G but also *t*ZOG were not affected in *ckx1* in contrast to wt where a substantial reduction occurred. Since O-glucosides CK species cannot be processed by CKXs, it appears that TOR inhibition specifically targets *t*Z N-glucosides which are known to be excellent substrates for CKX1. In this context, it is striking that *i*P9G is not affected since the affinity of CKX to this metabolite is even higher than for *t*Z9G (Galuszka et al., 2007), further underpinning the selectivity of TOR mediated effects towards *t*Z metabolites. Noteworthy, the strong accumulation of *c*Z metabolites in response to AZD8055 was the same as in wt for *ckx1* and *ckx1/3* (Fig. S17), suggesting that this effect is uncoupled from the effects on *t*Z metabolism.

**Figure 8:**
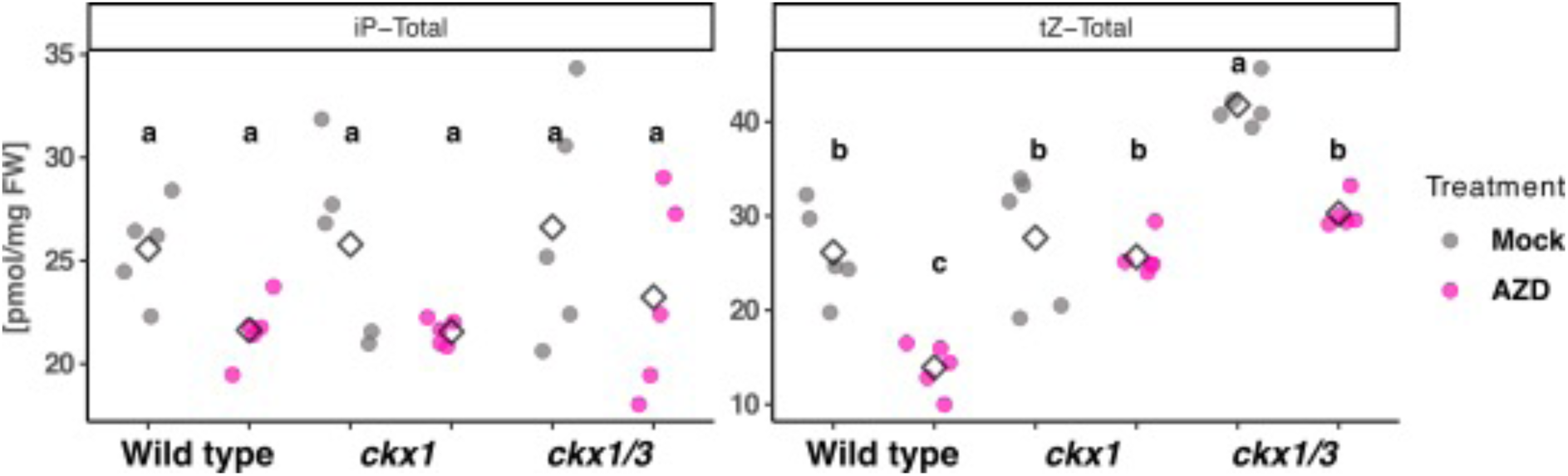
*ckx1* mutation rescues *tZ* deficiency caused by TOR inhibition. Quantification of different CK compounds in 4 day old seedlings treated with AZD8055 or mock for 8 h. *iP* = isopentenyladenine, *tZ* = *trans*-Zeatin. Letters indicate significance levels (p<0.05) obtained by linear model ANOVA with post-hoc t-test and bonferroni correction.

### CKX isoforms have specific effects on AZD8055 induced transcriptional reprogramming

Our physiological assays, together with the *pWUS* reporter assay as well as the CK profiling suggested a certain degree of redundancy or pleiotropy of different CKX isoforms in mediating changes of TOR activity. At the same time, not all *ckx* mutations conferred resistance to TOR inhibition and higher order mutants did not lead to additive phenotypes. Instead, the *ckx5/6* double mutant did not confer resistance to AZD8055 in our shoot growth assay while the individual single mutants both did. In addition, the *ckx5* mutation made the *pWUS* reporter resistant to TOR inhibition but the *ckx6* did not, clearly demonstrating specificity of individual CKXs. We hypothesized, that specific and redundant effects of different CKX isoforms should be resolved in diverging or overlapping transcriptional signatures of the respective mutant lines in response to TOR inhibition.

To address this experimentally, we sought to assess the effect of different single- and higher order *ckx* mutations on global TOR meditated transcriptional reprogramming. Thus, we performed another RNAseq experiment with *ckx1, ckx1/3, ckx1/3/5* (in Col-0 background, Fig.9 a + c), *ckx5, ckx6 ckx5/6* (in *pWUS:3xVenus:NLS/ pCLV3:mCherry:NLS* background, Fig.9 b + d) and their respective genetic background controls and evaluated their effects on AZD8055 induced transcriptional changes in whole seedlings. Similar to our previous experiment, TOR inhibition induced large scale transcriptional reprogramming with several thousand genes being differentially regulated in all genetic backgrounds (Table S2). The Venn - diagrams (Fig 9 a + b) reveal largely overlapping DEGs between the different mutants and their respective background control. However, at the same time a substantial proportion of DEGs is uniquely affected in specific *ckx* mutants, suggesting on the one hand that the core response is conserved, but on the other hand demonstrated that many transcriptional changes are exclusively dependent on specific CKX genes, which was in line with observations from other experiments. At the functional level, the top ten repressed GO categories enriched in the different *ckx* mutants were related to translation, protein biosynthesis and RNA processing in the wt (Fig. 9 c), as expected after TOR inhibition. Defense related GO terms, secondary metabolite biosynthesis and ABA response were enriched among the induced DEGs which again was in line with categories previously linked to TOR inhibition (P. Dong et al., 2015; Malinovsky et al., 2017; Scarpin et al., 2020; P. Wang et al., 2018). In *ckx1* and *ckx1/3* the GO terms among the repressed DEGs were very similar to the ones identified in wt. In contrast, we found no enrichment of GO terms related to translation or protein biosynthesis in the *ckx1/3/5* mutant and instead found defense and salicylic acid related terms to be highly represented among the repressed DEGs in this mutant. Among the induced GO categories, defense related terms were not found enriched in any of the mutant lines, indicating that TOR mediated defense response also depend on CKXs. For the *ckx6* and *ckx5/6* mutants, we again found GO terms related to translation mostly enriched among the repressed DEGs just as in the respective wt control (DR) (Fig. 9 d). Strikingly, the *ckx5* mutant stood out, as again none of the translation related terms was found, but instead terms related to “root development” were most prevalent. Conversely, “shoot system development” was the most enriched GO term in the *ckx5* mutant background. This nicely recapitulated our previous experimental observations where *ckx5* rescued shoot development on a global transcriptomic level. Again, defense related terms were absent from the induced GO categories in the *ckx* mutant lines. On the level of individual genes, we found several key regulators of stem cell fate and progression (e.g., ERECTA, MP, BAM3, CRN, PXY, REV, HEC2, UFO, STY1, ANT, YUC4 and others, see Table S3) specifically induced in the *ckx5* mutant, further demonstrating the central role of this specific isoform in TOR dependent control of stem cells. Additionally, a number of cell cycle regulators (CYCB2;3, CDKB2;2, CYCA1;1, ATE2F2, CYCB2;4, CDKB2;1, CYCA2;4, CYCB1;3, CYCD3;3, CYCD3;1, CYCB1;1, Table S3) was induced as well, which is surprising as TOR inhibition is thought to result in suppression of cell cycle. We also looked for differential regulation of *CKX* transcripts to elucidate cross regulatory or compensatory interactions. Indeed, we found *CKX1* expression to be induced in the *ckx5* and *ckx6* mutants. Similarly, *CKX4* RNA levels were strongly induced in the *ckx1/3* double mutant, but strongly repressed in the *ckx5* mutant. Hence, there appear to be complex, but highly specific regulatory interactions between CKX genes and the strong accumulation of *CKX4* may explain why *t*Z levels do still respond to AZD8055 in the *ckx1/3* mutans in the CK profiling experiment (Fig. 8).

**Figure 9:**
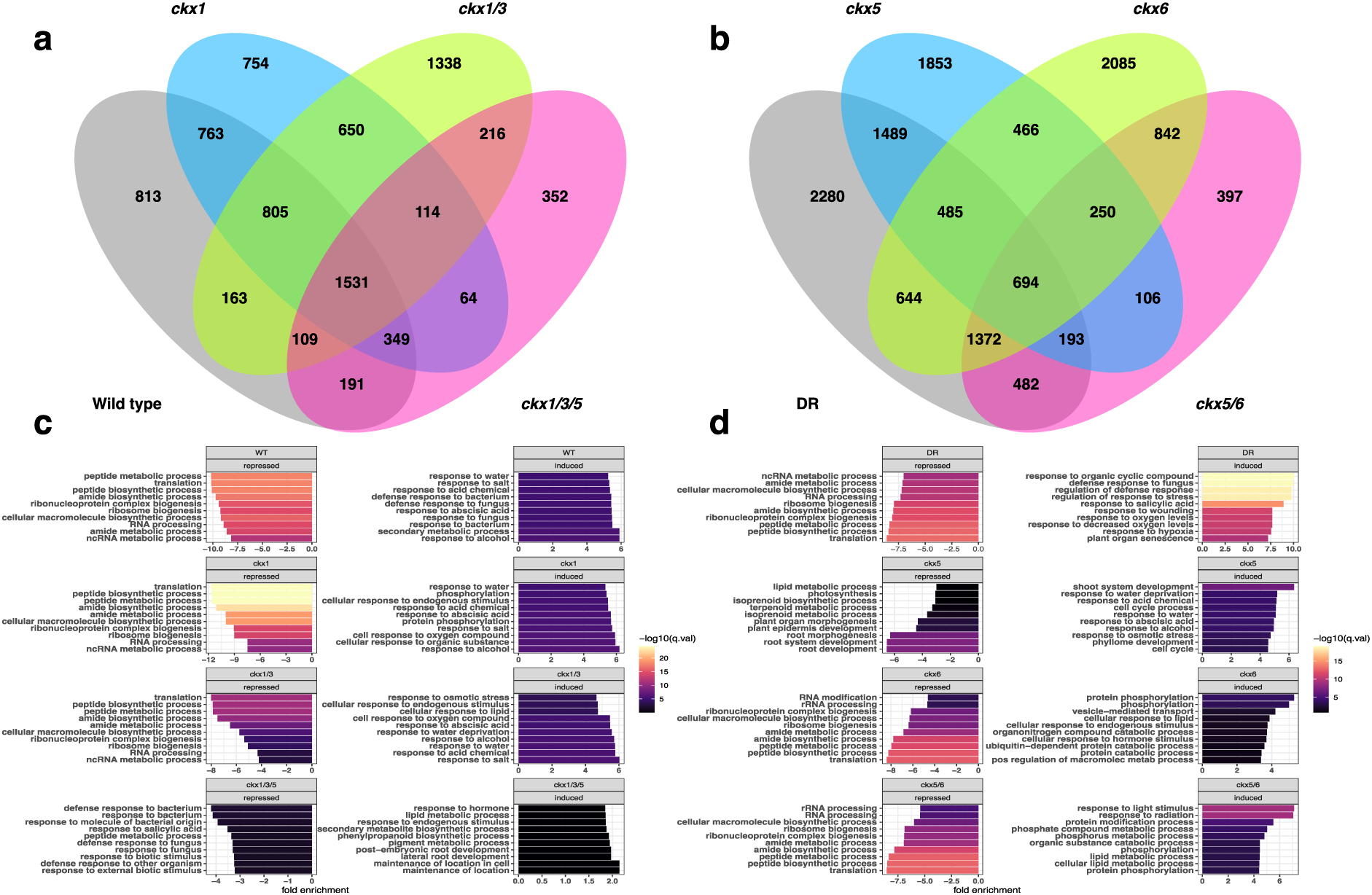
*ckx* mutations alter AZD8055 transcriptional profile. RNAseq data of whole seedling tissue from different *ckx* mutant lines treated with AZD8055 or mock for 8 h. DEGs obtained from DESeq2 analysis contrasting the effect of AZD8055 treatment in each genetic background. *ckx1*, *ckx1/3* and *ckx1/3/5* are in Col-0 background(Wild type). *ckx5*, *ckx6* and *ckx5/6* are in *pWUS:3xVenusNLS*, *pCLV3:mCherry:NLS* (DR) background. a) + b) Venn diagram of AZD8055 dependent DEGs in different *ckx* mutants. c) + d) Biological processes enriched in AZD8055 dependent DEGs in different *ckx* mutants.

Importantly, our transcriptional profiling confirmed that a core set of genes is affected by all CKX enzymes tested, but also that a substantial degree of isoform specificity exists. Particularly, CKX5 appeared to have a rather specific effect on the TOR dependent transcriptome, as in the *ckx5* single mutant but also in *ckx1/3/5* triple mutant the functional categories were drastically changed and translation, a core category of TOR function, was absent from the repressed DEGs. At the same time, genes related to shoot development were induced in the *ckx5* mutant pointing towards a central role of this isoform for TOR mediated control of shoot development, fitting well with the expression pattern of *CKX5* in SAM and vasculature (Bartrina et al., 2011).

### CK acts as upstream regulator of TOR

The observation that translation did not appear as repressed GO category after TOR inhibition in the ckx5 and ckx1/3/5 mutants suggested that CK might also act as an upstream regulator of TOR activity. We therefore treated etiolated seedlings with BA and monitored S6K phosphorylation as readout for TOR activation (Fig. 10). In line with our hypothesis, BA treatment led to strong induction of S6K phosphorylation which suggested that CK indeed is able to activate TOR. As previously shown, ROP2 mediates auxin dependent activation of TOR, hence we were interested if ROP2 might also be involved in CK mediated TOR activation. Indeed, the induction of S6K phosphorylation after BA treatment was almost completely abolished in the *rop2-1* mutant or after induction of a dominant negative ROP2 construct (DN-ROP2, D120A). This demonstrated that CK acts as an upstream regulator of TOR in a ROP2 dependent manner similar to auxin.

**Figure 10:**
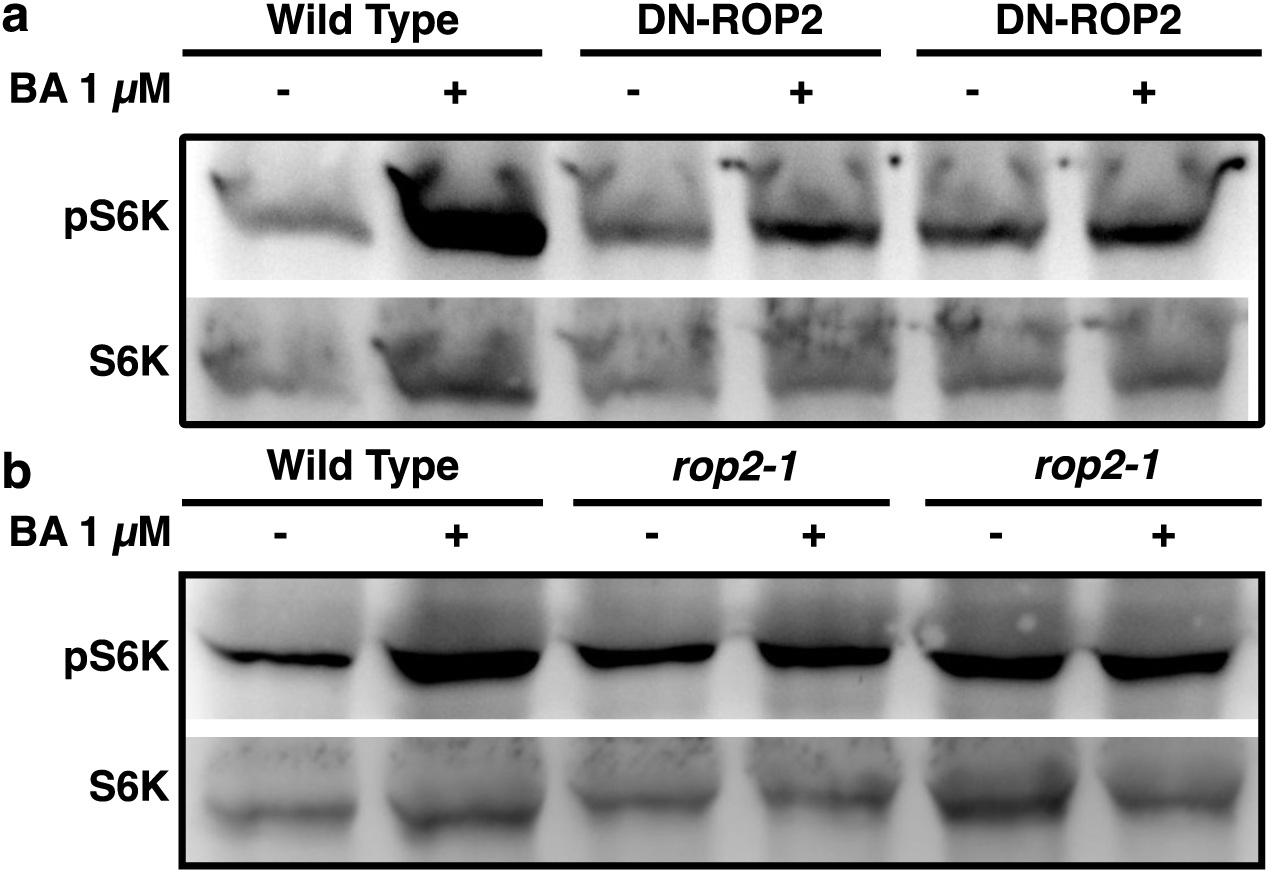
Cytokinin activates TOR in a ROP2 dependent manner. a) + b) Western blots of protein extracts form 3 d old etiolated seedlings. Membranes were probed with serum detecting TOR specific phosphor-epitope (T449) or total protein of S6K1/2. Treatment with benzyladenine (BA) for 24 h. a) All seedlings were treated with 10 *µ*M estradiol for induction of dominant negative ROP2 (D120A) (DN-ROP2) expression. *rop2-1* SALK_055328C.

In sum, our results demonstrated that the TOR kinase controls *WUS* expression and partially shoot development via regulation of cytokinin levels, specifically *tZ*. This effect is mediated through modulation of protein accumulation of specific CKX cytokinin degrading enzymes. Mechanistically, TOR controls transcription of several CK pathway genes, and restricts CKX protein accumulation, by repressing *CKX* mRNA translation. Moreover, CK also acts as an upstream regulator of TOR activity which puts TOR in a feed forward loop where TOR controls CK levels which in turn activate TOR.

## Discussion

Plants dynamically adjust their transcriptional and translational programs to quickly adapt to changing environments. The TOR kinase network is a master regulator of cellular metabolism and controls translation in accordance with nutrient availability in all eukaryotes. Several reports have shown a connection between TOR and stem cell activity in plants and identified upstream regulators for this specific function (Li et al., 2017; Pfeiffer et al., 2016; Xiong et al., 2013). However, the downstream effectors connecting TOR with stem cell regulatory programs so far remained elusive. Focusing on shoot stem cells, we show that TOR controls accumulation of specific CK enzymes by translational repression of their mRNAs (Fig. 6a, Fig. 7 + S15). CKX enzyme activity in turn limits the availability of *trans*-zeatin, the major instructive signal for SAM development and *WUS* expression (Fig. 3+4+5b+c).

CKX enzymes catalyze the irreversible degradation of CKs and hence exert strong influence on development and growth (Bartrina et al., 2011). CKs are potent regulators of plant growth and development and act as long-distance signaling molecules that instruct cellular identity and behavior of diverse tissues, including the SAM (Kieber & Schaller, 2014; Landrein et al., 2018; Osugi et al., 2017). Thus, controlling CK abundance is key to synchronizing SAM activity with cellular homeostasis of distant source tissues that take up or produce the resources necessary for organogenesis and growth. Our results suggest that TOR under favorable conditions represses translation of CKX mRNAs, whereas when TOR is inhibited CKX translational efficiency is enhanced resulting in globally reduced levels of *trans*-zeatin and ultimately in reduced activity of stem cells in the SAM. We hypothesize that this active translational repression enables global adjustments of CK availability faster compared with transcriptional adjustments and may therefore grant a fitness advantage.

Intriguingly, not all CK metabolites appear equally affected by TOR inhibition as *iP* levels are largely unaffected while most *tZ* metabolites are reduced. Similarly, CK response towards exogenously added *tZ* is impaired while response towards *iP* is not, both suggesting differential degradation of specific CK metabolites. This is further corroborated by the complete rescue of TOR mediated *t*Z depletion in the *ckx1* mutant. The simplest explanation are different selectivity profiles of CKX enzymes towards certain CK derivatives, however, the fact that TOR inhibition seems to majorly act on *t*Z-glucosides but CKXs have an even higher affinity *in vitro* towards *i*P-glucosides argues against this notion (Galuszka et al., 2007; Kowalska et al., 2010). Instead, differential compartmentalization of different CKs and CKXs provides a coherent model that could explain our observations, even more so, as previous evidence suggests differential subcellular localization of different CK species (Jiskrová et al., 2016).

Our results did not allow us to draw conclusions about the detailed molecular mechanism by which TOR activity results in translational repression of specific transcripts. However, one potential mechanism might involve SnRK1 dependent translation mediated by eIFiso4G, although in published translational efficiency data of eIFiso4G mutants no evidence for differential translation of CKX isoforms was found so far (Cho et al., 2019). Alternatively, GCN2 dependent translation via eiF2alpha could drive translation antagonistically to TOR (Y. Dong et al., 2017; Sesma et al., 2017).

Our polysome data showed very specific responses of mRNAs encoding specific CKX enzymes in seedling shoots, raising the question of regulatory specificity. While diverse CKX transcripts exhibit structural differences that may underly their specific responses to TOR dependent translational repression, differential regulatory effects may also arise from different cellular contexts. Cell type specific TOR functions have been suggested by a recent study showing differences in TOR activity between source and sink tissues resulting in inverse regulation of plasmodesmata permeability (Brunkard et al., 2020). Since we have carried out polysome profiling on entire seedling shoots, but CKX transcripts have highly specific expression pattern (Bartrina et al., 2011; Werner et al., 2003), some of the differences we have observed may therefore come from divergent tissue context, which we could not resolve. Some of our observations appeared inconsistent between different experiments for specific *ckx* mutants. Particularly, the inverse behavior of *ckx6* and *ckx5/6* mutants between the physiological- and *WUS* reporter assays was puzzling. The *ckx6* mutant showed some resistance in the shoot growth assay even though we observed no differences on the polysome level, albeit *CKX6* was the only transcript that showed mild accumulation in qRT-PCR after TOR inhibition which could explain how CKX6 is linked to TOR. However, the *ckx6* mutation did not affect *WUS* response towards TOR inhibition, which could be due to expression pattern allowing to affect one CK dependent trait but not the other. Along these lines, also the behavior of *ckx5/6* double mutants is surprising, as they did show an additive effect in the *WUS* reporter assay but were not resistant in the shoot growth assay even though both single mutants were. The transcriptome profiles of the *ckx* mutants suggested some antagonistic relationship between *ckx5* and *ckx6*. *ckx5* single mutants exhibited very different functional profiles in its list of DEGs after TOR inhibition compared to *ckx6* mutant seedlings. For example, translation was not overrepresented among the GO categories in *ckx5*, whereas shoot development specifically was identified among the DEGs with increased expression. Surprisingly, in *ckx5/6* double mutants the enriched gene sets were similar to wt and translation was identified as major functional category among repressed DEGs. At this point we can only speculate that differences in the expression domains and/or substrate specificity between CKX5 and CKX6 may drive specific functions and that hormonal imbalance between tissues may result in complex regulatory effects in single mutants. The relationship of *ckx5* to *ckx1/3* appears much simpler as the GO categories of the *ckx1/3/5* triple mutants resemble *ckx5* single mutants.

Our work suggests that control of *t*Z catabolism is the dominant mechanism regulating stem cells in the shoot and shoot growth downstream of TOR. However, our data also show that there is no simple linear relationship between TOR and CKXs, but that instead both are intertwined in a complex feedback relationship that very likely depends on the tissue context. This exemplifies that shoot development is a complex trait controlled by intricate genetic interactions affecting multiple physiological and molecular processes. While previous studies do show a correlation between *tZ* abundance *WUS* expression and shoot growth, our data reveals that *WUS* expression and thus stem cells only represent a part of the shoot growth program. Importantly, it is evident that TOR controls a large diversity of other growth-related processes such as cellular expansion, proliferation and cellular homeostasis also supported by the finding that several thousand genes respond to its inhibition.

Interfering with CK homeostasis through targeted expression of CKXs can increase plants resilience to environmental stresses such as drought or cold and improve yield traits (Ashikari et al., 2005; Cortleven et al., 2019; Nehnevajova et al., 2019; Schwarz et al., 2020; Werner et al., 2010). Thus, CKX enzymes represent promising candidates of agronomic importance (Jameson & Song, 2020). Consequently, modulating their translational control bears the potential to generate cultivars that ameliorate the detrimental effects of broad CKX overexpression. Along these lines, we observed that the supplementation of sugars together with CKs is sufficient to drive *WUS* expression even when TOR is inactive. This further fuels the question about specific TOR functions in stem cells compared to differentiated tissues, since in this context CKs are not necessarily degraded in the SAM, although some CKX isoforms are specifically expressed in the OC (Bartrina et al., 2011; Werner et al., 2003).

Taken together, our findings on TOR mediated translational repression of catabolic enzymes open new avenues to study the mechanisms of TOR driven environmental adaptation and stem cell control. Considering the ecological benefits that could conceptually be derived from tunable translational repression of growth factor catabolism, this regulatory logic may be much more widespread than anticipated so far. Thus, future studies might reveal analogous mechanisms for other catabolic genes in plants or animal systems.

## Materials and methods

### Plant material

All used plant lines were in the Col-0 background. The double reporter line pWUS:3xVenus:NLS/ pCLV3:mCherry:NLS as well as the *ckx5*, *ckx6* and *ckx5/ckx6* CRISPR mutants are described in (Pfeiffer et al., 2016). The p35S:ARR1ΔDDK:GR line was described in (Sakai et al., 2001) and was crossed with the double reporter line to obtain homozygous alleles for each transgene. The p35S:cMyc-CKX1 line is described in (Niemann et al., 2015, 2018). The ckx2 (SALK_083761c), ckx3 (SALK_050938c), ckx4 (SALK_055204c) mutants were obtained from NASC. The *ckx1* mutant is an EMS allele recovered from the tetraploid Arabidopsis TILLING population (Tsai et al., 2013)that contains a G-to-A transition at nucleotide 1252 of the *CKX1* gene (At2g41510)leading to a premature stop codon (Trp311*). *ckx3* and *ckx5* mutant alleles were described by (Bartrina et al., 2011).

The *cre1-2/ahk3-7* mutant was described in (Riefler et al., 2006). The rop2-1 and the DN-ROP2 line have been described in (Denninger et al., 2019)

### Growth conditions

Seeds were sterilized with 70% ethanol and 0.1% Triton for 10 min and afterwards washed twice with autoclaved water. Seeds were plated on 100 *µ*m nylon meshes (nitex 03/100–44, Sefar, Heiden, Switzerland) on top of 0.5x MS (Duchefa, Haarlem, The Netherlands), 0.9% Phytoagar in square petri dishes. After plating, seeds were imbibed for 3 days at 4°C in darkness and transferred to growth cabinets (poly klima, PK 520-LED, Freising, Germany) where they were kept under continuous light at 22°C and grown vertically for 4 days. Subsequently, seedlings were transferred with the nylon meshes to 0.5x MS plates supplemented with 2 *µ*M AZD8055 (Selleckchem, Houston, TX) or equal volumes of DMSO for 8 h.

### Liquid culture

About 30–40 seeds, that were imbibed as described above, were sown in 3 ml 0.5x MS in petri dishes of 35 mm diameter. Plants were kept in darkness for three days after the induction of germination by 6 hr light treatment. The medium of two day old etiolated seedlings was supplemented with the indicated treatments. All stock solutions were 1000x concentrated and diluted in DMSO, therefore control plants were mock treated with the same volume of DMSO.

### RNAseq

Seedlings were grown as described under growth conditions. 4 days after germination seedlings were transferred on a mesh to 0.5xMS plates containing either 2 *µ*M AZD8055, 10 *µ*M TORIN1, 20 *µ*M KU63794 or equal volumes of DMSO as mock control. After 8 h, 30 mg of shoot tissue were harvested for each replicate and frozen in liquid nitrogen. 3 independent replicates were harvested for each condition. Total RNA was extracted with the Plant RNA Purification Reagent (Invitrogen, Carlsbad, CA) according to the instructions of the manufacturer, digested with TURBO DNAse (Ambion/ Thermo Fisher, Waltham, MA) and purified with RNeasy Mini Kit (Quiagen, Hilden, Germany). Libraries were poly-(A) selected and analyzed with NEXTseq 500. For differential gene expression analysis reads were aligned with RNA STAR (v2.6) alignment tool with TAIR10 genome model as reference. Transcripts were assembled and counted with StringTie (v1.3.3) and statistical analysis was performed using DESeq2 (v1.18.1) (Love et al., 2014). GO term analysis was performed using ThaleMine web tool (https://www.bar.utoronto.ca/thalemine/begin.do).

The second RNAseq experiment was performed as the first one. Reads were aligned using SALMON and differential expression using DESeq2 (v1.18.1). GO-term enrichment analysis has been performed using the “gage” package based on a KEGG database.

### Histochemical GUS staining

Four day old seedlings were harvested in 90% acetone and incubated at −20°C for at least 1 hr. Seedlings were washed with PBS and incubated in substrate buffer (1x PBS (pH 7.0), 1 mM K_3_Fe(III)(CN)_6_, 0.5 mM K_4_Fe(II)(CN)_6_, 1 mM EDTA, 1% Triton X-100, 1 mg/ml X-gluc) at 22°C over night. After staining, the seedlings were incubated with 60% and subsequently in 95% ethanol to remove chlorophyll.

### Microscopy and fluorescence quantification

All images were obtained using Zeiss Imager M1, the Plan-APOCHROMAT 20x/0.8 objective (Zeiss, Oberkochen, Germany) and YFP- and GFP-specific filter sets. Procedures for fluorescent reporter activities of the double reporter were performed as described in (Pfeiffer et al., 2016). Each measurement was normalized to the median (set to 100) of the mock for experiments in the light or to the 6-BA treated samples for experiments performed in the dark.

### Western blot

Approximately 30 mg of shoot tissue were harvested, and proteins were extracted with 1:4 ratio (mg/*µ*l) adjusted to the exact fresh weight with 95°C hot denaturing buffer (100 mM MOPS pH 7.6, 100 mM NaCl, 40 mM ß-MeOH, 5% SDS, 10% Glycerol, 4 mM EDTA, 2 mM PMSF, PI (Sigma)) and boiled at 95°C for 5 min. Cellular debris was removed by two centrifugation steps (10 min, 14,000 rpm, RT). Equal volumes of the obtained extract were separated on a 10% SDS-PAGE gel and blotted to a PVDF membrane. Membranes were probed with Phospho-p70 S6 kinase (Thr(P)-389) polyclonal antibody (No.9205, Cell Signaling Technology, Cambridge, UK) to detect S6K phosphorylation. S6K1/2 antibody (AS12-1855, Agrisera AB, Vännäs, Sweden) was used to detect total S6K1 and S6K2. c-Myc antibody (9E10, Santa Cruz Biotechnology, Dallas, TX) was used to detect c-Myc tagged CKX1.

### Physiology

Seeds were singled out on 0.5xMS, 0.9% Phytoagar plates and imbibed for three days at 4°C in the dark. Plates were kept horizontally in long day conditions at 22°C for four days. *t* 40 single seedlings at the same developmental stage and of similar size were selected and transferred to plates containing the indicated AZD8055 concentrations and grown for seven more days before shoot fresh weight was measured. For the measurements, seedling shoots were removed and weighed in batches of 5 – 10 seedlings. Afterwards the average weight was calculated for each batch.

### Statistical testing

Statistical analysis for experiments shown in Fig. 5a-e, linear models were (with “R”) were generated with Genotype (Freshweight *t* AZD concentration*Genotype) analyzed with ANOVA and post-hoc t-test to calculate significance levels. Datasets were previously evaluated for extreme outliers, normality assumptions and heteroscedasticity. Pairwise t-tests have been performed for group comparisons with Bonferroni correction. Statistical analysis was performed in R (v4.0.2) with unnormalized data. ED50 values were calculated using the “drc” package for dose response analysis in R (Ritz et al., 2015). The dose of effect ratios in Fig. S12 were calculated with the *EDcomp* function from the “drc” package with the *delta* method to estimate confidence intervals (Ritz et al., 2015).

Data that was not normally distributed was tested with Wilcoxon rank test and Hochberg correction. Normally distributed data was tested for heteroscedasticity and two-tailed students t-test with equal or unequal variance have been performed accordingly.

### CHX chase assay

P35S:cMyc-CKX1 seedlings were grown as described under growth conditions. 8 h after transfer to 2 *µ*M AZD8055 the plates were flooded with 50 ml 200 *µ*M cycloheximide and 2*µ*M AZD8055 solution (0.015% Silwet L-77) for 0, 2, 4 and 8 h and shoots were harvested for western blot analysis as described above.

### RT-qPCR

Total RNA was extracted as described under RNAseq. RNA integrity was confirmed on an agarose gel and the concentrations were determined with a nanodrop device. Equal amounts of RNA were used for oligo dT primed cDNA synthesis with the RevertAid First Strand cDNA Synthesis Kit (Thermo Fisher, Waltham, MA). The qPCR reaction was set up using the SG qPCR Master Mix (EURx, Gdansk, Poland) and run on a qTOWER^3^ (Analytic Jena, Jena, Germany) PCR System with technical duplicates each.

### Cytokinin profiling – LC-MS

For cytokinin profiling seedlings were grown as described for RNAseq analysis and 5 biological replicates of shoot tissue were harvested for each condition. The CK content was determined by ultra-high performance liquid chromatography-electrospray tandem mass spectrometry (Svačinová et al., 2012), including modifications described by (Antoniadi et al., 2015). Briefly, samples (20 mg FW) were homogenized and extracted in 1 ml of modified Bieleski buffer (60% methanol, 10% HCOOH and 30% H_2_O) together with a cocktail of stable isotope-labeled internal standards (0.25 pmol of CK bases, ribosides, *N*-glucosides, and 0.5 pmol of CK *O*- glucosides, nucleotides per sample added). The extracts were purified onto an Oasis MCX column (30 mg/1 ml, Waters) and then analyzed using using an Acquity I-class system (Waters, Milford, MA, USA) combined with a mass spectrometer Xevo™ TQ-XS (Waters, Manchester, UK). Data were processed with Target Lynx V4.2 software and final concentration levels of phytohormones were calculated using isotope dilution method (Novák et al., 2008).

### Polysome fractionation

200 mg plant material grown as described under growth conditions was homogenized by rotating at 4°C in 600 *µ*l polysome extraction buffer (0.2 mM Tris-HCL, pH=9, 0.2 mM KCL, 25 mM EGTA, 35 mM MgCl2, 1% DOC, 1% PTE, 1% Brij-35, 1% Triton X-100, 1% NP-40, 1% Tween-20, 5 mM DTT, 10 *µ*M MG-132, 50 *µ*g/ml Cycloheximide, 50 *µ*g/ml chloramphenicol and 1% EDTA-free protease inhibitor cocktail). Extracts were centrifuged at 16000xg at 4°C for 10 min. 300 *µ*l supernatant was loaded to 7-47% sucrose gradient and centrifuged at 38000x rpm for 3 hours in a Beckmann SW41Ti rotor. The gradient was fractionated after recording the absorbance at 254 nm. RNA was precipitated from 1 ml fraction by mix and incubation with one volume of 8 M guanidine-HCL and two volumes of absolute ethanol at - 20°C over night followed by centrifuge at max. speed for 1 hour. RNA pellet was resuspended with 50 *µ*l DEPC water. 100 ng RNA was used for cDNA synthesis (SuperScript IV reverse transcriptase (ThermoFisher, 18090050) which was subsequently analyzed by qRT-PCR as described above.

### Cytokinin response assay

Seedlings were grown as described under growth conditions. After 8 h of AZD8055 or mock treatment seedlings were sprayed with an atomizer with either 100 nM of trans-zeatin (Duchefa, Haarlem, The Netherlands) or 100 nM of isopentenyladenine (Duchefa, Haarlem, The Netherlands) solution (0.015% Silwet L-77). After 30 min three independent replicates of shoots and roots were harvested separately for total RNA extraction and RT-qPCR analysis as described above.

## Supporting information

Table of differentially expressed genes

## Acknowledgements

We thank Sebastian Wolf and Aurelio Teleman for their ideas and scientific input. The authors give sincere thanks to Hana Martínková and Petra Amakorová for their help with phytohormone analyses. This work was supported by the ERC grants #282139 “StemCellAdapt”, and #810296 “DECODE” and the SFB873 (DFG) to JL, as well as the Ministry of Education, Youth and Sports of the Czech Republic (European Regional Development Fund-Project “Plants as a tool for sustainable global development” No. CZ.02.1.01/0.0/0.0/16_019/0000827).

## Author contributions

O.N. and M.S. performed the cytokinin profiling experiment. Y.D. performed the ribosome fractionation experiment and gave input for results interpretation, L.A.R. gave input for results interpretation. I.B. and T.W. generated the ckx1 and derived higher order mutants and gave input for results interpretation. A.P. performed one WUS reporter assay generated the cross between the DR line and p35S ARR1∂DDK:GR and designed experiments, D.J. performed all other experiments, designed experiments, curated and analyzed the data, wrote the first draft, revised and edited the manuscript, J.U.L. designed experiments, edited the manuscript and acquired the funding.

