## Supplementary material for "TOR kinase controls shoot development by translational repression of cytokinin catabolic enzymes": Supplemantary Information

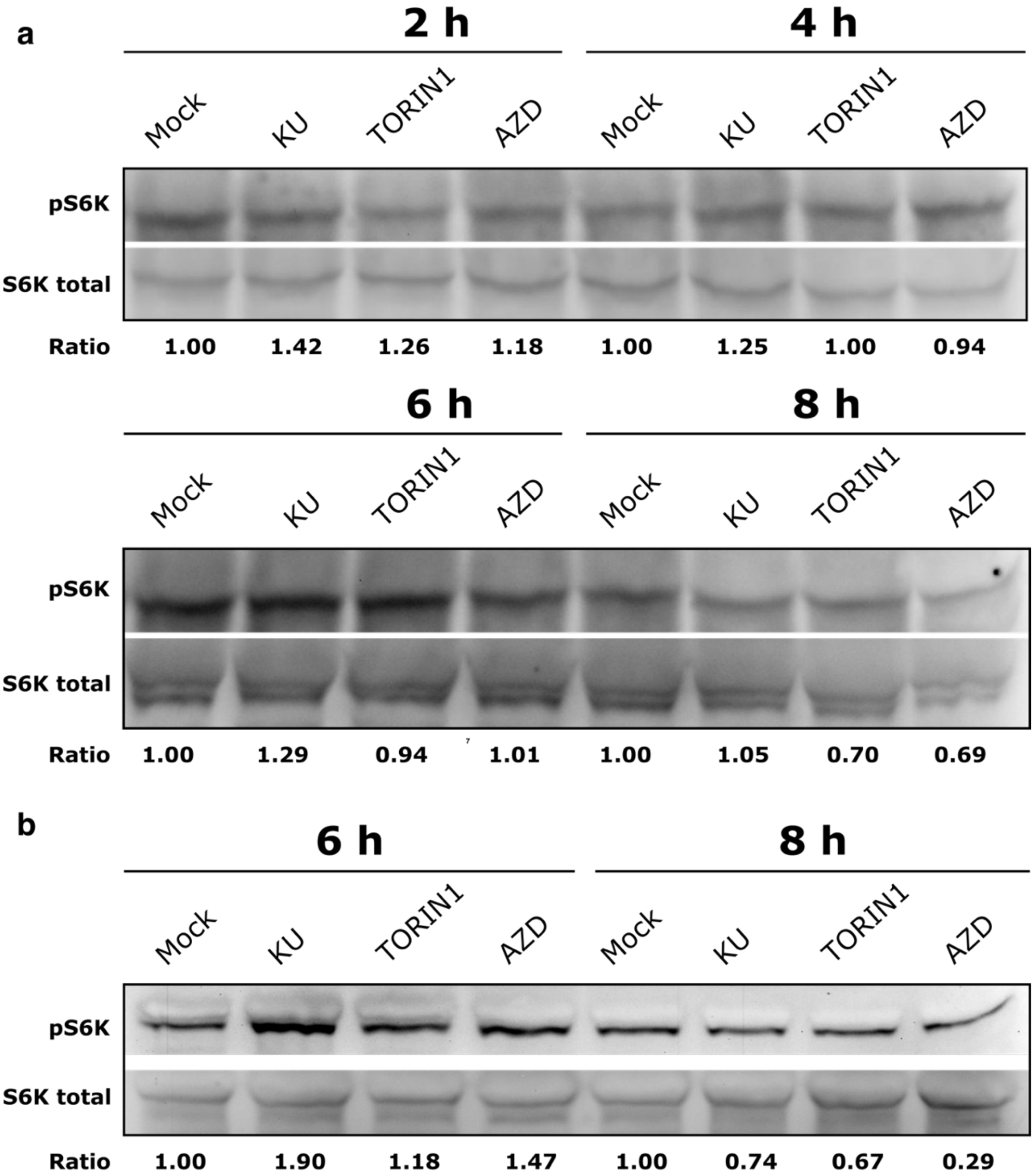


**Figure S1: TOR activity drops 8 h after transfer**

Western blot of Arabidopsis shoot tissue extracts from 4-day old seedlings transferred to plates containing different TOR inhibitors for the indicated time. Membranes were probed with serum detecting TOR specific phosphor-epitope (T449) or total protein of S6K1/2.

**a)** Concentrations: 2 $\mu$M AZD8055, 10 $\mu$M TORIN1, 10 $\mu$M KU63794. Ratios were calculated between pS6K (No.9205, Cell Signaling) and S6K total band intensities and normalized to the respective Mock sample which was set to 1.

**b)** Concentrations: 2 $\mu$M AZD8055, 10 $\mu$M TORIN1, 20 $\mu$M KU63794. Ratios were calculated between pS6K (abcam, ab207399) and S6K total band intensities and normalized to the respective Mock sample which was set to 1.

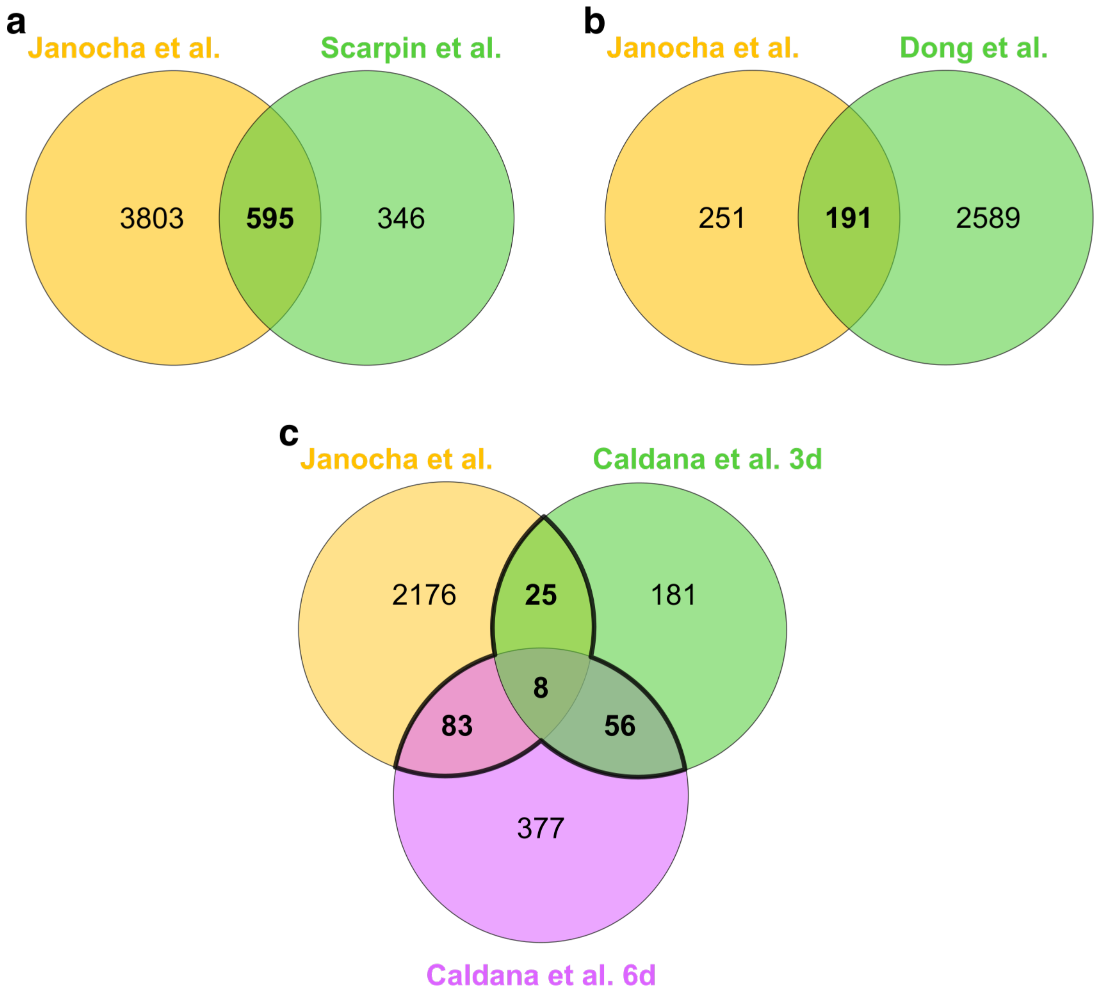


**Figure S2: RNAseq data overlaps with other TOR related datasets**

- **a)** Venn diagram of DEGs from this study compared with TORIN2 regulated DEGs from (Scarpin et al., 2020). Hypergeometric test reveals 4.5 fold enrichment of the overlap with p = 2.6 e-276.
- **b)** Venn diagram of DEGs from this study with compared with AZD8055 regulated DEGs from (Dong et al., 2015). Hypergeometric test reveals 4.86 fold enrichment of the overlap with p = 8.86 e-84. Expression cutoff is log2 fold change > 1.
- **c)** Venn diagram of DEGs from this study compared with DEGs of inducible *tor* RNAi lines from (Caldana et al., 2013) after 3d of induction (green) or 6d of induction (purple). Hypergeometric test reveals 1.22 fold enrichment which is not significant (p = 0.3) after 3d and significant (p = 1.07 e-07) 1.74 fold enrichment after 6d of *tor* RNAi induction. Expression cutoff is log2 fold change > 0.5.

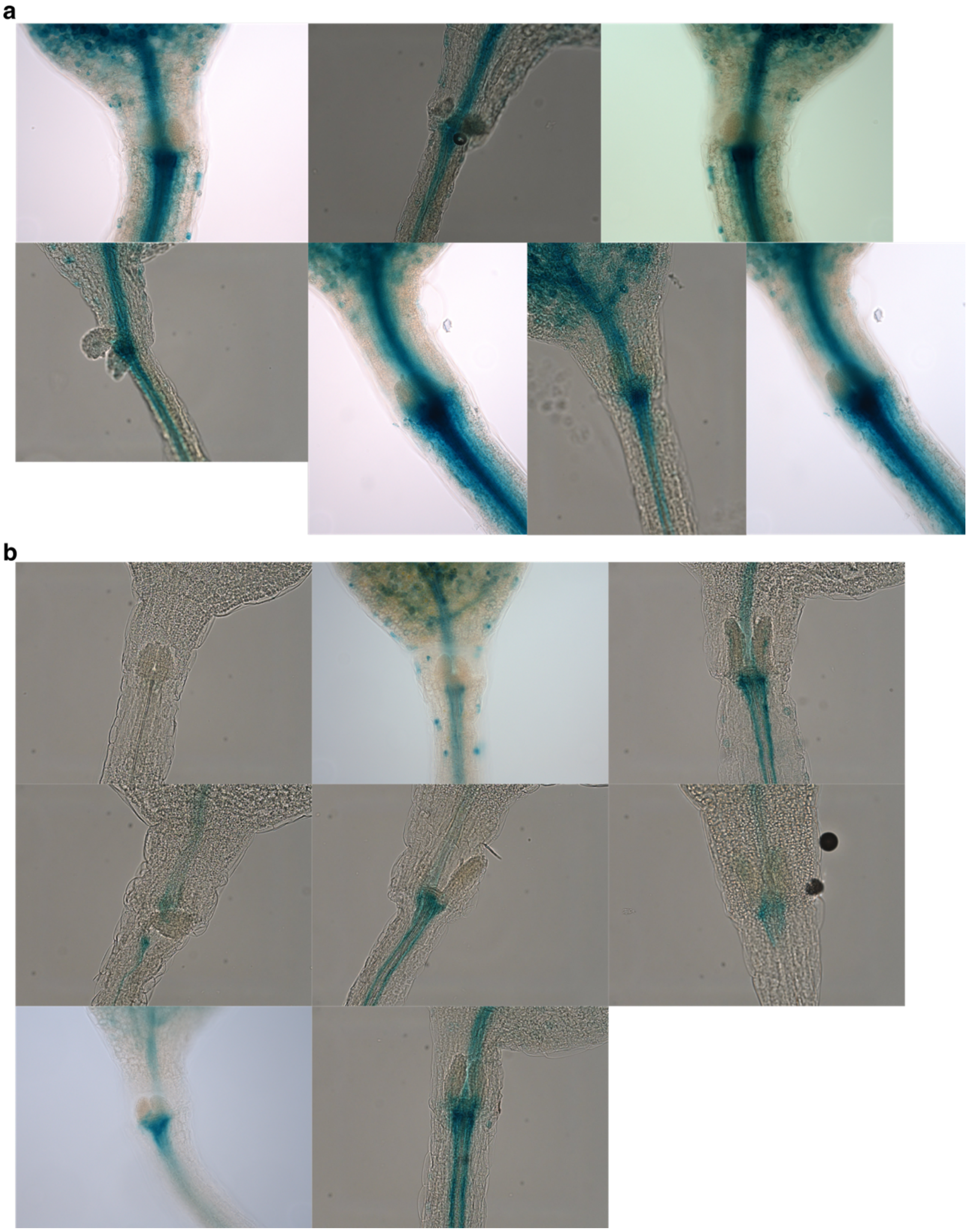


**Figure S3: Cytokinin signaling in the SAM is reduced upon TOR inhibition**
**a)+b)** Microscopic images of pTCSn:GUS reporter treated for 24 h with a) Mock or b) 2*µ* AZD8055.

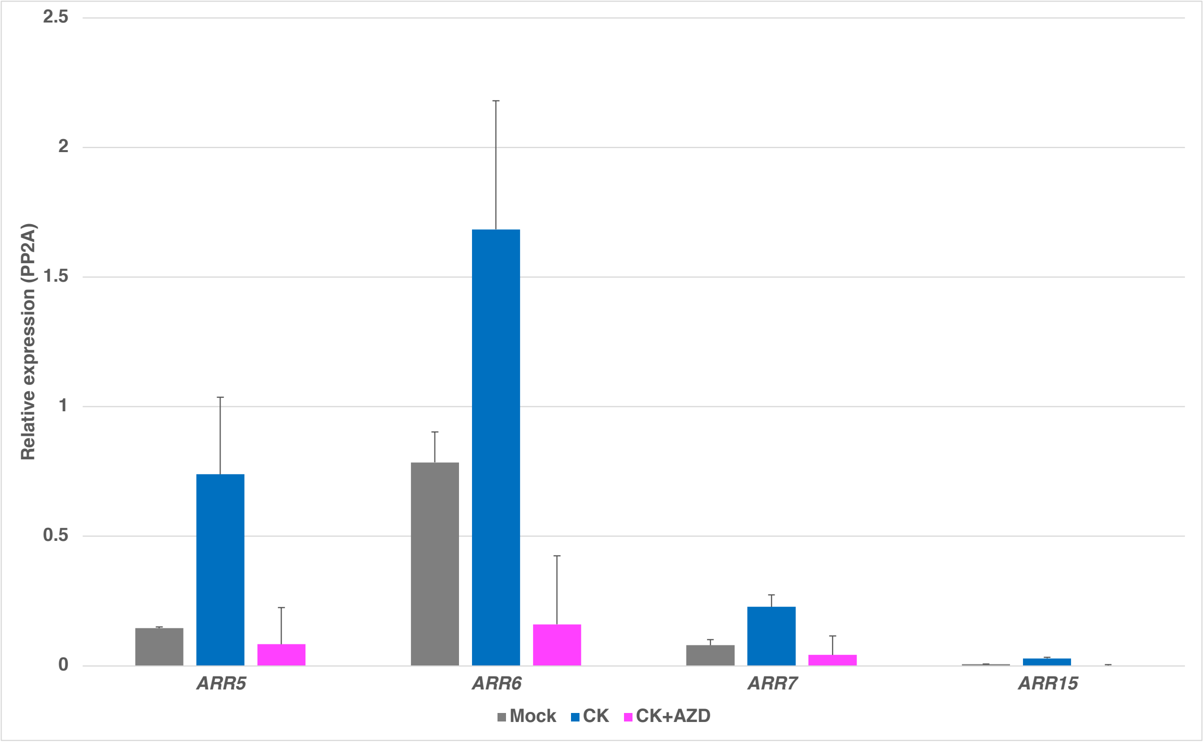


**Figure S4: TOR inhibition reduces CK signaling output**

q-RT-PCR of 3 day old etiolated seedling shoots grown for 3 days on mock, 0.5 $\mu$M 6-BA or 6-BA + 2 $\mu$M AZD8055 supplemented medium. Error bars represent standard deviation of 3 biological replicates.

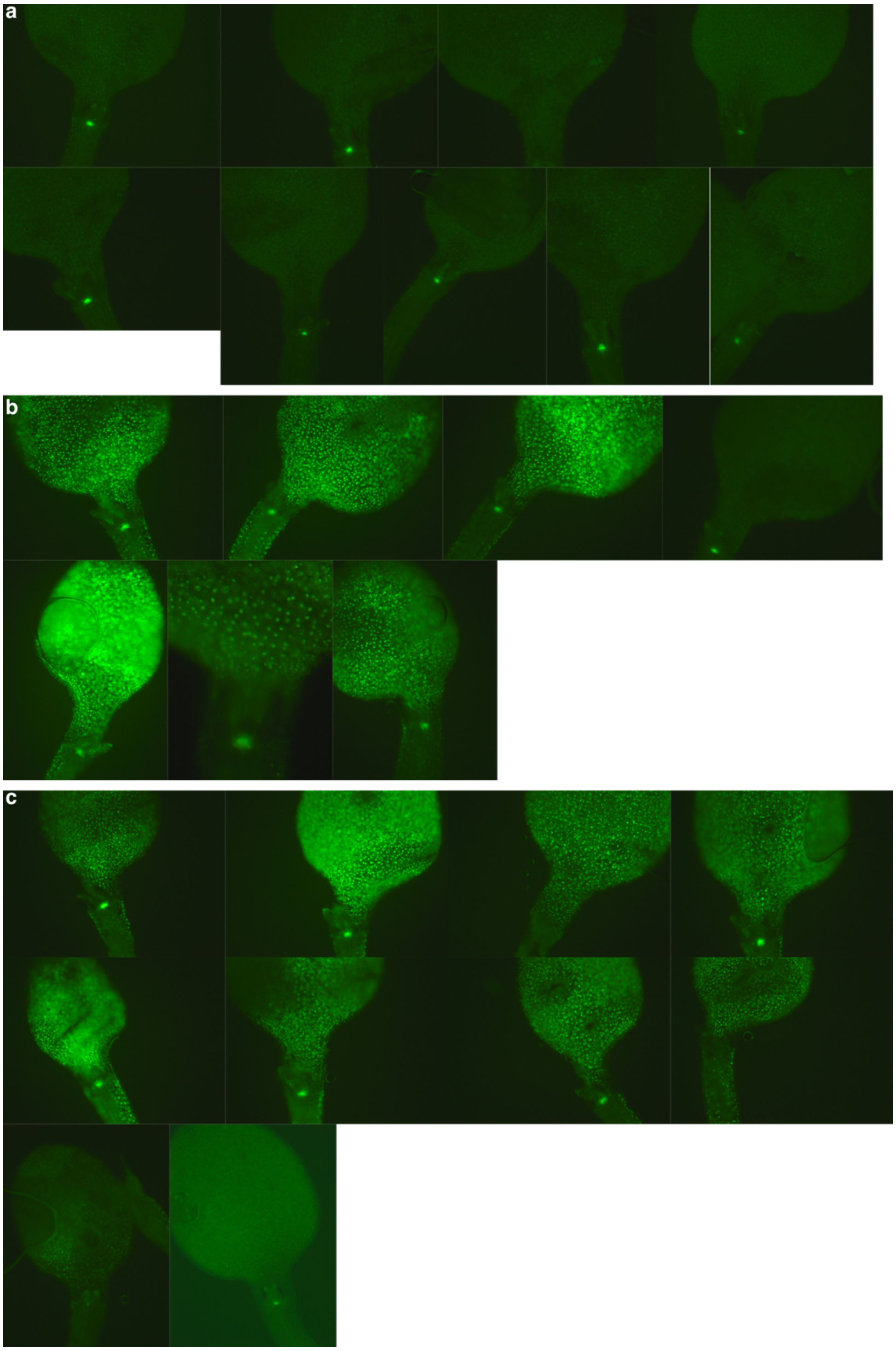


**Figure S5: TOR does not affect ARR1∂DDK activity**

**a)-c)** Microscopic images of pWUS:3xVenus-NLS reporter line crossed with p35S:ARR1∂DDK-GR. 4 day old seedlings treated with a) mock, b)10 $\mu$M dexamethasone (DEX) or c) DEX + 2$\mu$M AZD8055. Refers to Figure 2b-d.

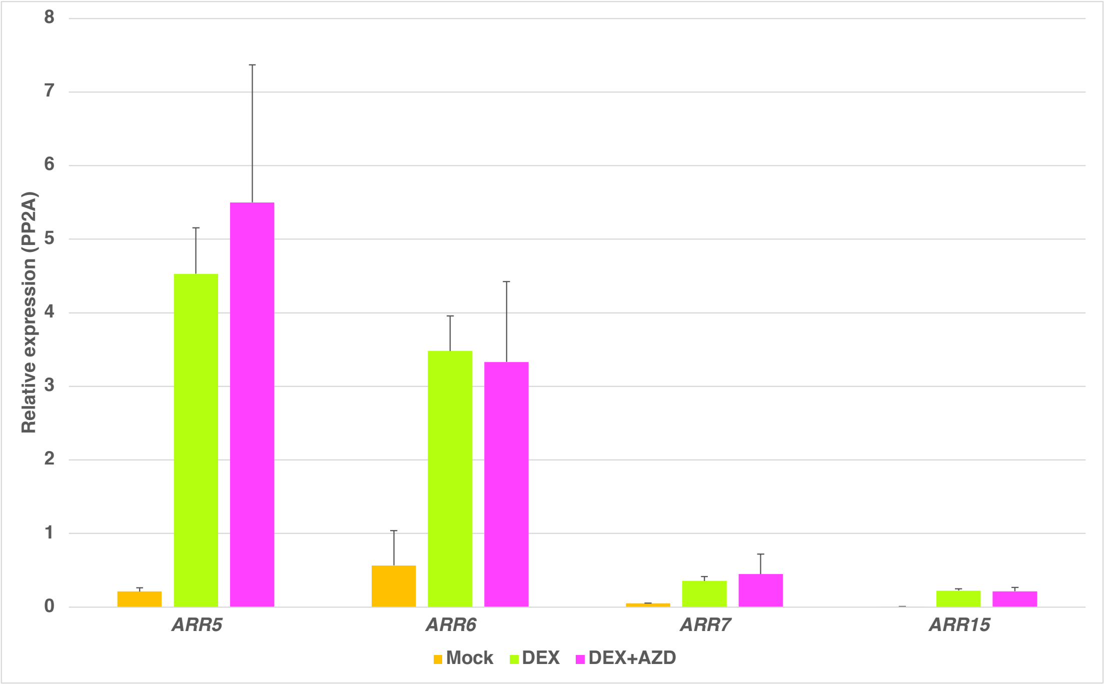


**Figure S6: ARR1∂DDK induced CK signaling is TOR independent**

q-RT-PCR of 3 day old shoots grown for 1 day on either Mock, 10 $\mu$M dexamethasone (DEX) or DEX + 2 $\mu$M AZD8055. Error bars represent standard deviation of 3 biological replicates.

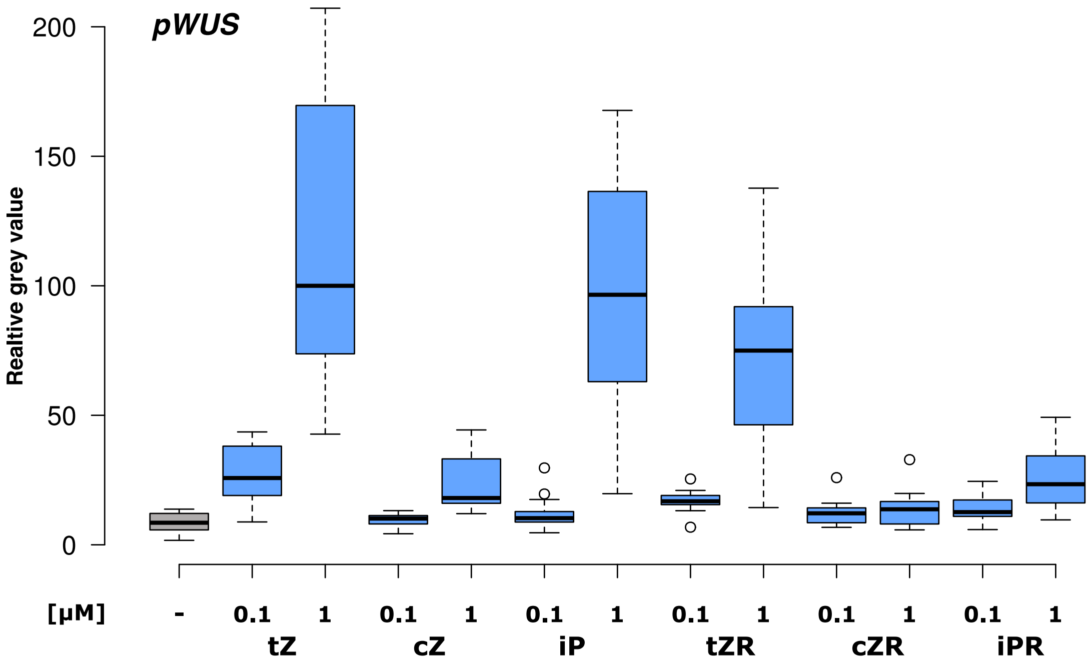


**Figure S7: Potential of different CKs to induce *pWUS***

Quantification of pWUS:3xVenus:NLS reporter signal. 2 day old etiolated seedlings were treated with mock or the indicated concentrations of different CK derivatives for 3 days.

*t*Z = *trans*-Zeatin, *c*Z = *cis*-Zeatin, *i*P = *iso*pentenyladenine, *t*ZR = *trans*-Zeatin riboside, *c*ZR = *cis*-Zeatin riboside, *i*PR = *iso*pentenyladenosine.

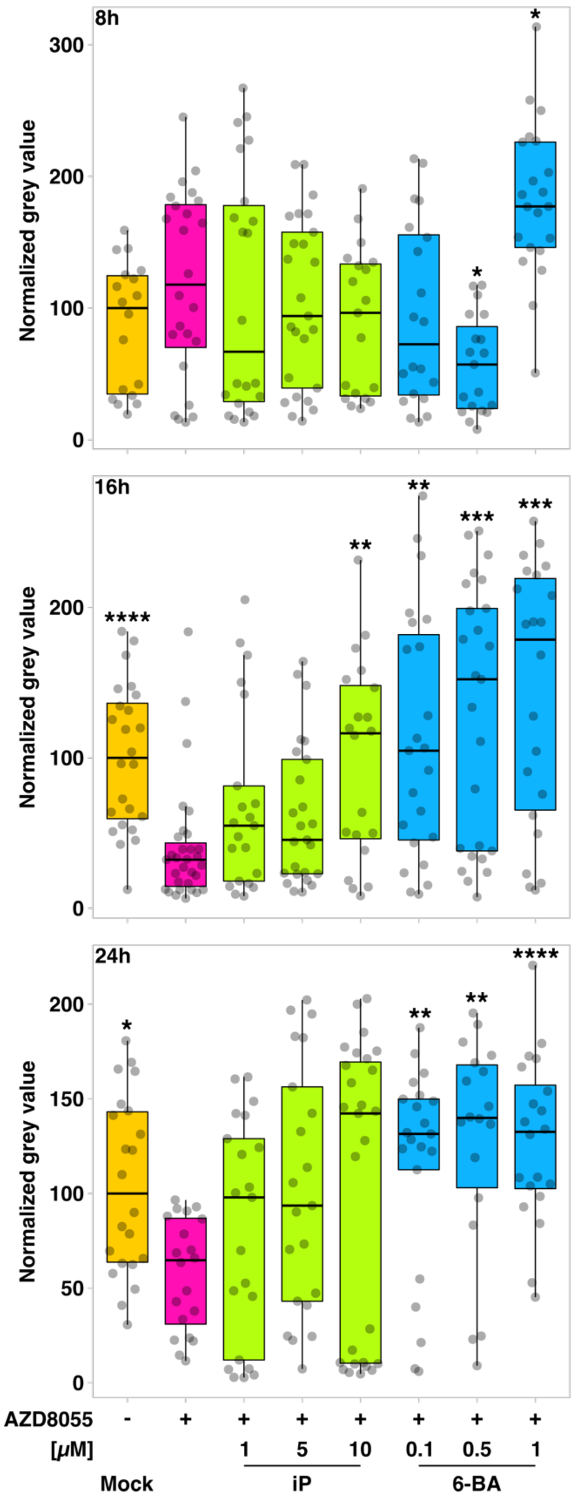


**Figure S8: Cytokinins rescue WUS expression upon TOR inhibition**

4 day old pWUS:3xVenus-NLS seedlings were treated for the indicated time with mock or 2 $\mu$M AZD8055 and the indicated cytokinin concentrations. Grey dots represent single measurements. Asterisks indicate significant differences to the respective AZD treated condition calculated with Wilcoxon rank sum test with Hochberg correction (* = $p < 0.05$; ** = $p < 0.01$; *** = $p < 0.001$; **** = $p < 0,0001$). iP = isopentenyladenine; 6-BA = 6-benzylaminopurine.

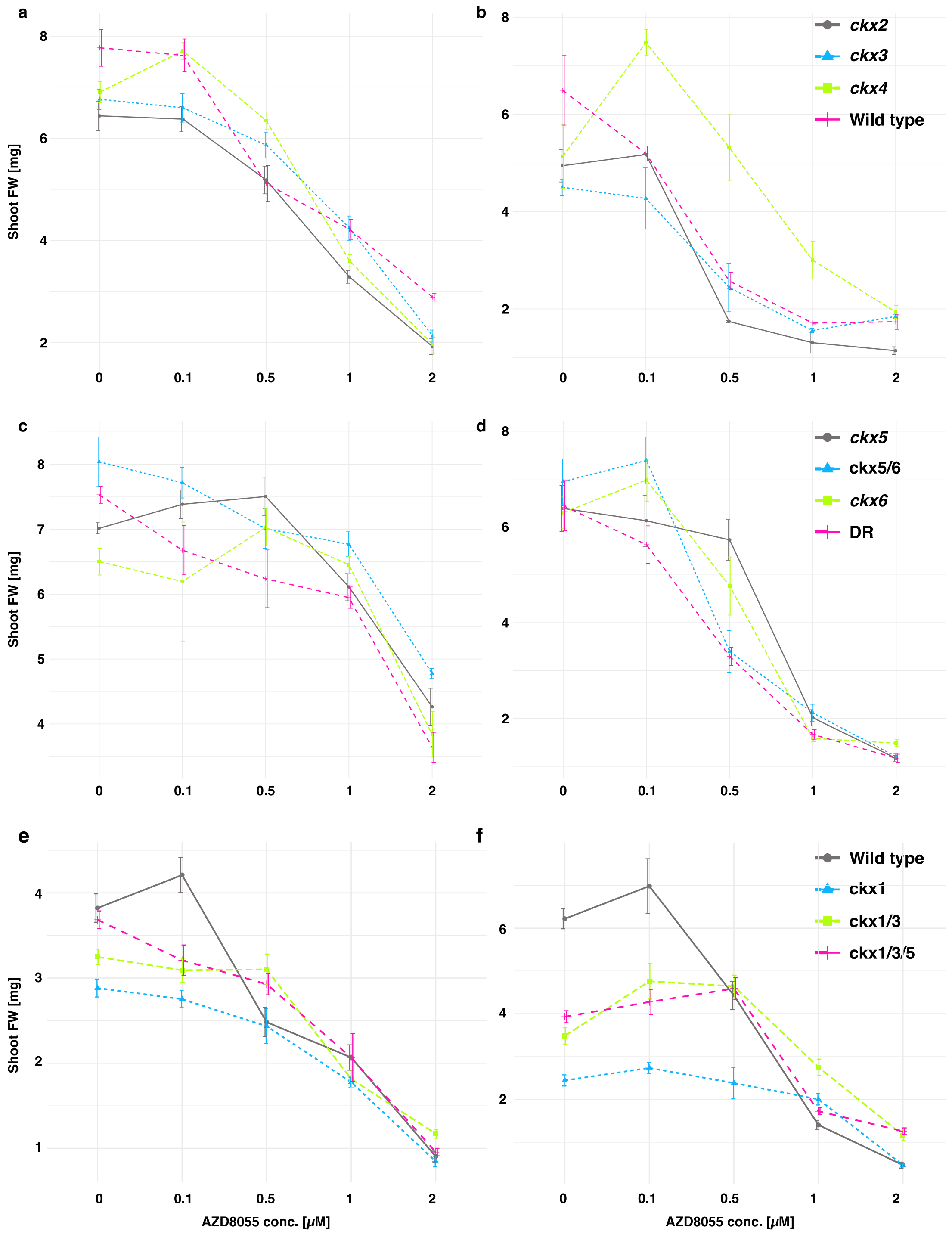


**Figure S9: Dose response curves of *ckx* mutants**

Line plots showing single experiments data from Fig. 5a+b+c before normalization.

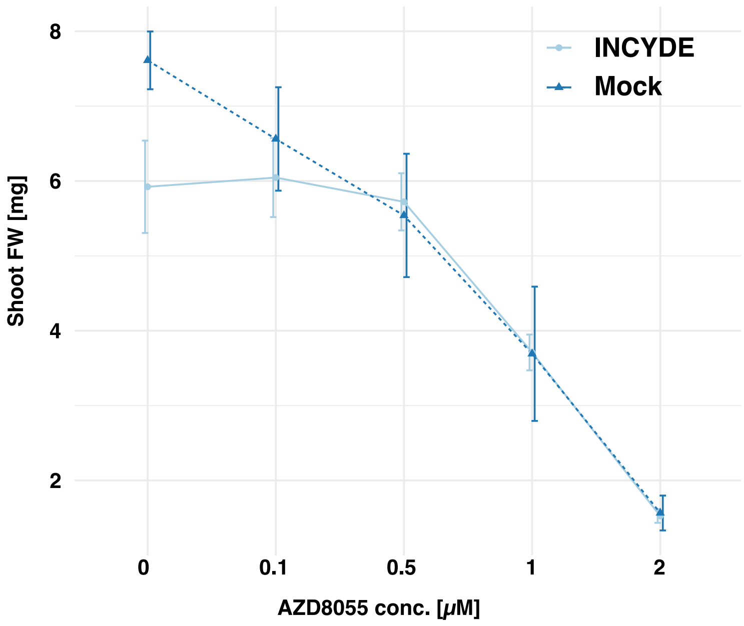


**Figure S10: Dose response curves of INCYDE treatment**

Line plots showing single experiments data from Fig. 5e before normalization.

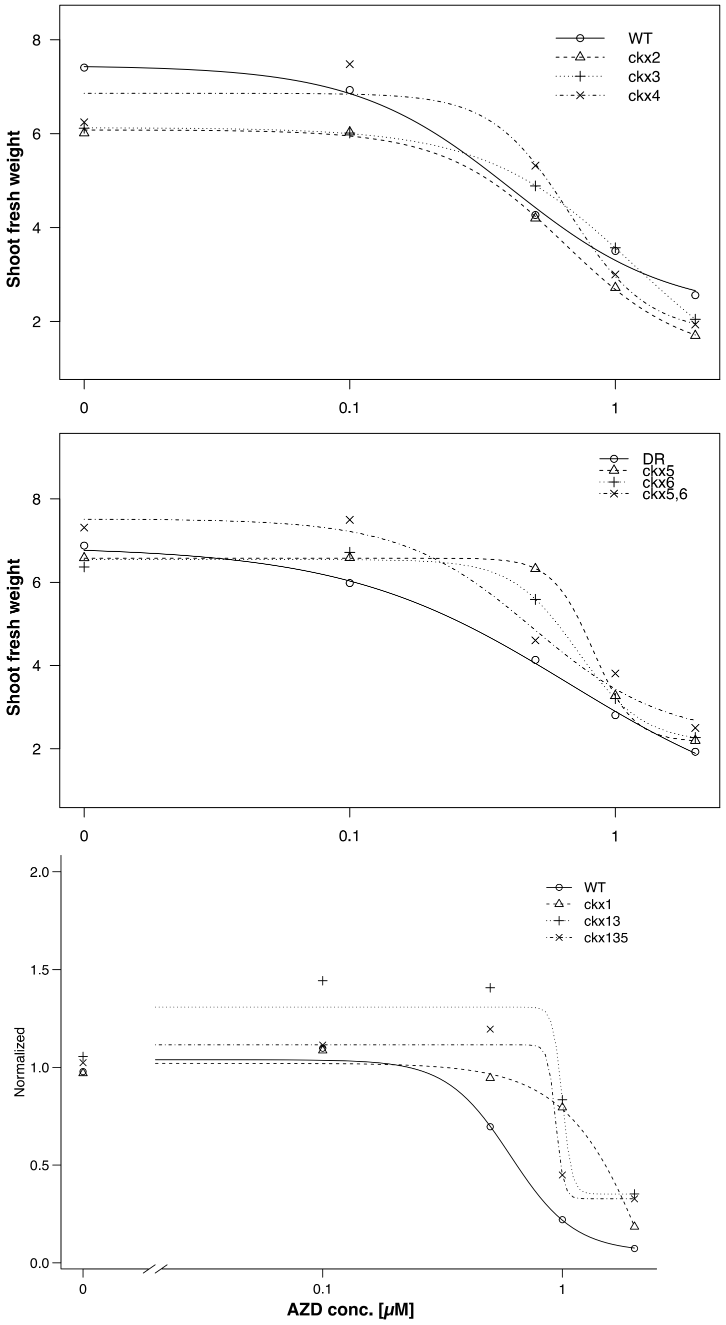


**Figure S11: Dose response curve model fit**

Model fit for the physiological data from Fig5a+e obtained by the “drc” package from “R”. Data before normalization was used to generate the statistical models.

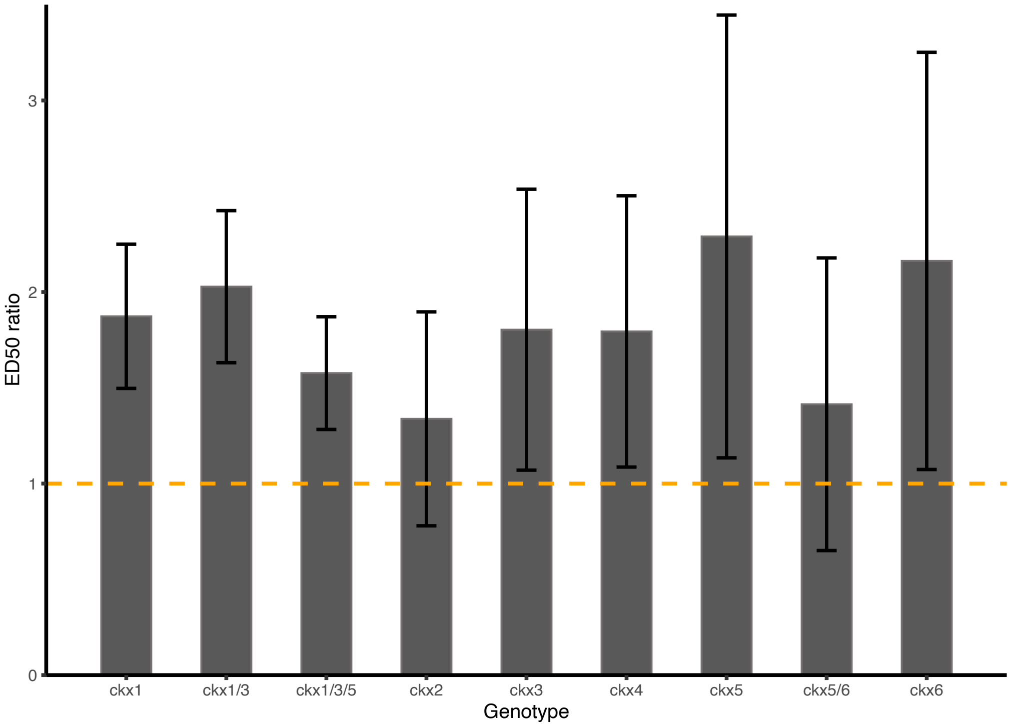


**Figure S12: Dose of effect values of different *ckx* mutant lines**

Bar plots show AZD8055 [$\mu$M] dose of effect ratios for 50% growth inhibition for different genotypes relative to wild type (indicated by orange dashed line). Values were determined from shoot fresh weight data from Figure 5a+e using the “drc” package in “R”. Error bars represent 95% confidence intervals.

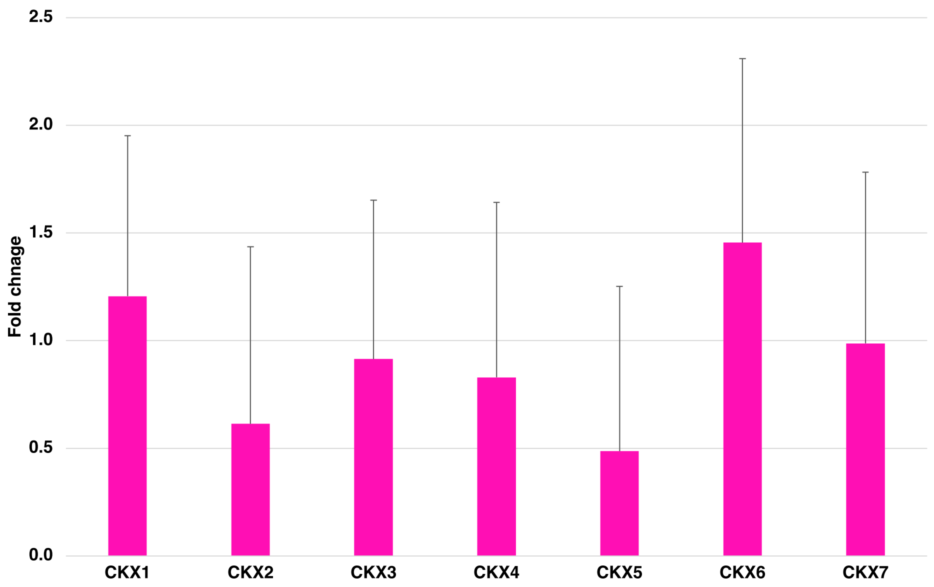


**Figure S13: CKX transcriptional changes upon TOR inhibition**

q-RT-PCR of RNA from the RNAseq experiment. Expression was normalized to PP2A expression and the fold change of the AZD8055 treated samples was calculated relative to the mock. Error bars represent confidence interval of 3 biological replicates. Students t-test reveals significant differences for *CKX2* (p = 0.015), *CKX5* (p =0.031) and *CKX6* (p = 0.045).

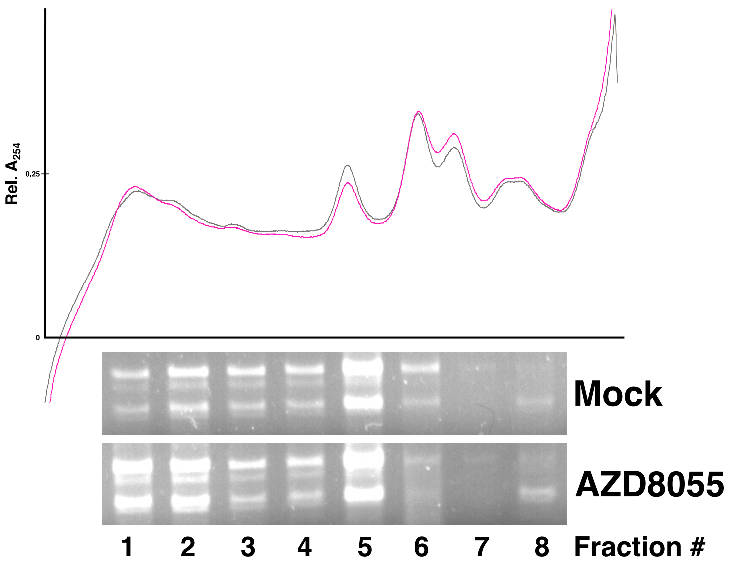


**Figure S14: Ribosome fractionation**

Representative absorption spectrum and RNA gel blot of the corresponding fractions.

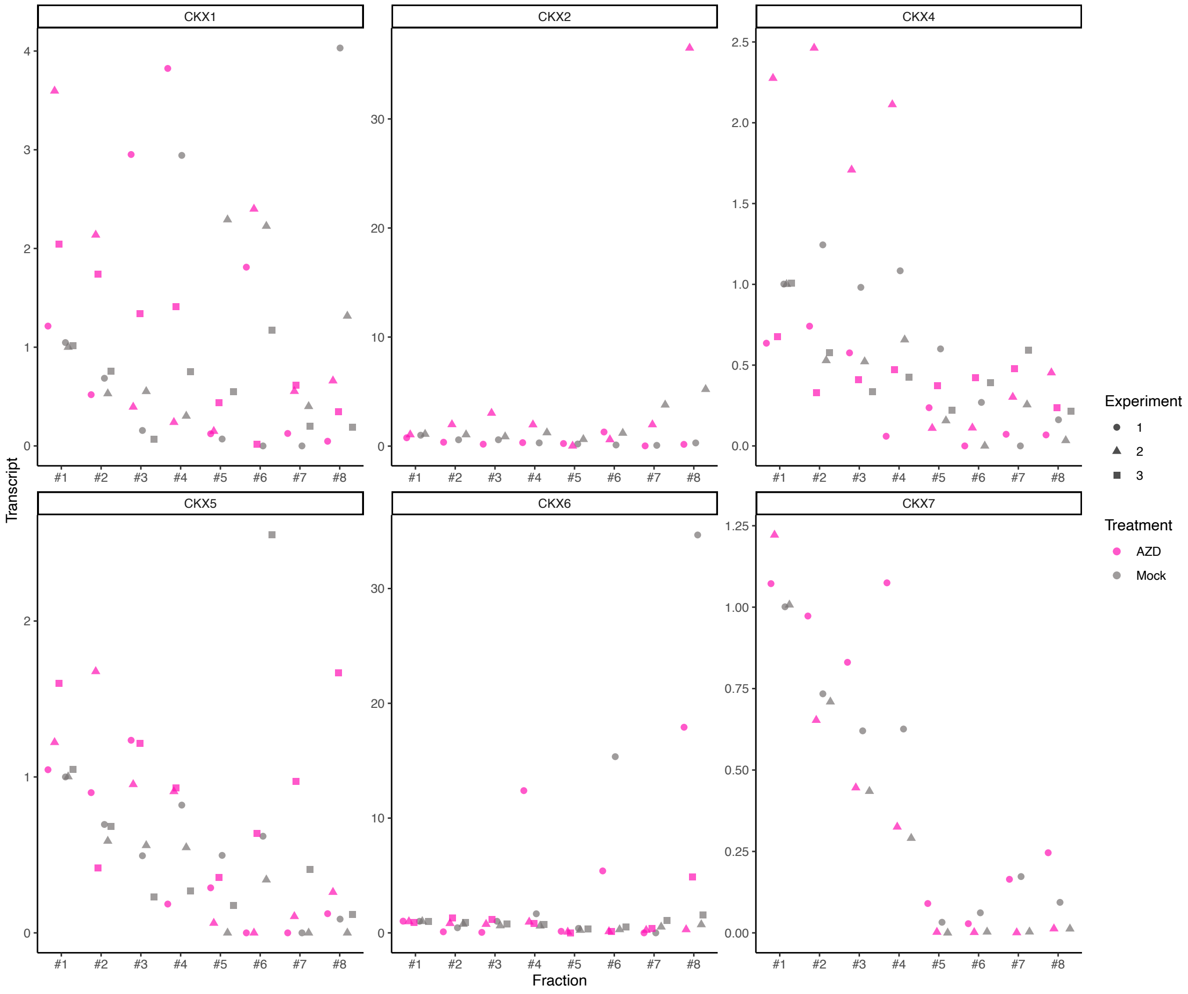


**Figure S15: TOR dependent translational regulation of CKX isoforms**

Ribosome fractionation experiments. Heavy polysomal fractions (fractions 1-3), light polysomal fractions (fractions 4+5) and monosomal fractions(fractions 6-8).CKX transcripts were detected with q-RT-PCR relative to *UBI10* transcript and normalized to the respective mock of fraction #1. Datapoints correspond to independent experimental repetitions corresponding to Figure 7.

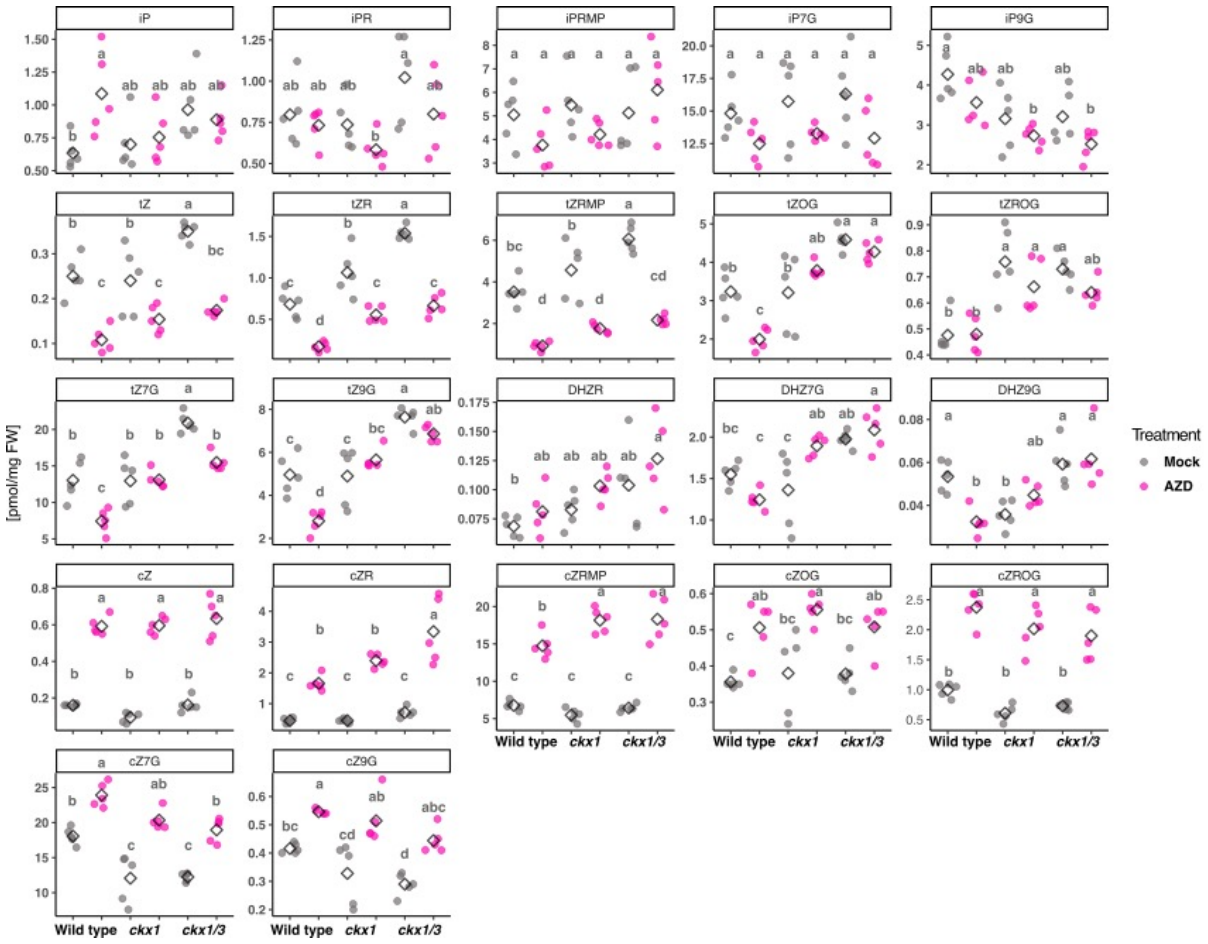


**Figure S16: CK metabolic profiling**

LC-MS profiling of different CK metabolites. Letters indicate significance levels obtained by linear model ANOVA with post-hoc t-test and Bonferroni correction.

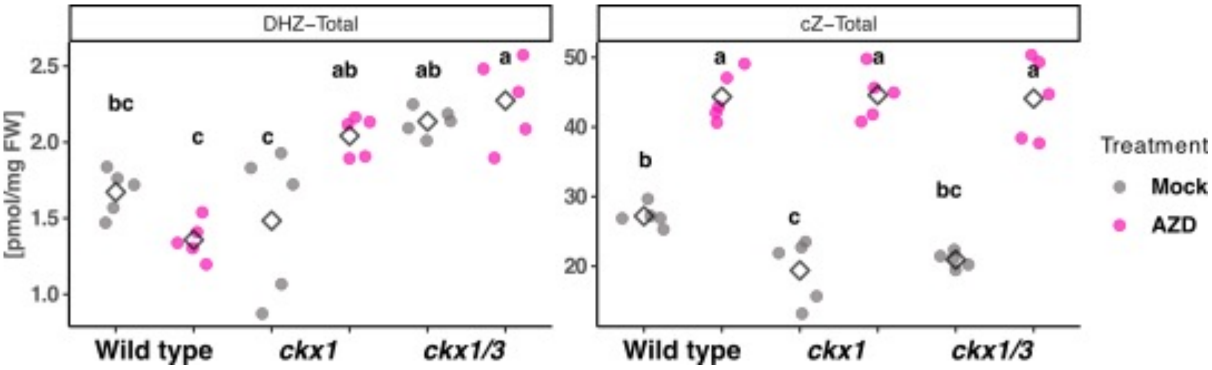


**Figure S17: CK metabolic profiling**

LC-MS profiling of different CK metabolites. Letters indicate significance levels obtained by linear model ANOVA with post-hoc t-test and Bonferroni correction.

**Primer List for q-RT-PCR**

| ID | Primer Name | Sequence |
|---|---|---|
| A08633 | CKX7fwd_qPCR | CGGAGTCAATGGTCCAATGC |
| A08634 | CKX7rev_qPCR | GAACCGAAGCAATGCCACAA |
| A08635 | CKX6fwd_qPCR | CCCAGTCATCGTCTACCCAG |
| A08636 | CKX6rev_qPCR | CGATGTTAGGATCGCCACCA |
| A08637 | CKX5fwd_qPCR | GTTCCAACGGCTCTGTTTTGT |
| A08638 | CKX5rev_qPCR | CCGTTGTAAAGACCGATGTCG |
| A08639 | CKX4fwd_qPCR | ATAACGAGGGCCAGGATTGC |
| A08640 | CKX4rev_qPCR | AGTCAACTCCGAGATCATTGGT |
| A08641 | CKX3fwd_qPCR | ACCGCGAAGAAAAGATCCGA |
| A08642 | CKX3rev_qPCR | AACGGCGGAATTAGTGGACA |
| A08643 | CKX2fwd_qPCR | CTCTGGTATCATCGCCGACA |
| A08644 | CKX2rev_qPCR | CTTCGGGACTCGCTCTTCTC |
| A08645 | CKX1fwd_qPCR | TTCCACACAGGCAAGCAGAT |
| A08646 | CKX1rev_qPCR | ACTTGCCAGTTTCCTGATCCAT |
| A09146 | mycCKX1 fwd | AGACTTGAACGGACTCGACG |
| A09179 | mycCKX1 rev | CGAGGAAAGTCTTGTTGTTT |
| A01067 | PP2A_fwd | TAACGTGGCCAAAATGATGC |
| A01068 | PP2A_rev | GTTCTCCACAACCGCTTGGT |

Primers used for type-A ARRs were described in (Zhao et al., 2010).
Primers used for normalization of polysome data AT4G05320 UBQ10 fwd: GGCCTTGTATAATCCCTGATGAATAAG rev: AAAGAGATAACAGGAACGGAAACATAGT
